# Novel environment exposure drives temporally defined and region-specific chromatin accessibility and gene expression changes in the hippocampus

**DOI:** 10.1101/2024.10.31.621351

**Authors:** Lisa Traunmüller, Erin E. Duffy, Hanqing Liu, Stella Sanalidou, Elena G. Assad, Senmiao Sun, Naeem S. Pajarillo, Nancy Niu, Eric C. Griffith, Michael E. Greenberg

## Abstract

The interaction of mammals with a novel environment (NE) results in structural and functional changes in multiple brain areas, including the hippocampus. This experience-dependent circuit reorganization is driven in part by changes in gene expression however, the dynamic sensory experience-driven chromatin states and the diverse cell type specific gene expression programs that are regulated by novel experiences are not well described. We employed single-nucleus multiomics (snRNA- and ATAC-seq) and bulk RNA-seq of the hippocampal DG, CA3, and CA1 regions to characterize the temporal evolution of cell-type-specific chromatin accessibility and gene expression changes that occur in 14 different cell types of the hippocampus upon exposure of mice to a novel environment. We observe strong hippocampal regional specificity in excitatory neuron chromatin accessibility and gene expression as well as great diversity in the inhibitory neuron and non-neuronal transcriptional responses. The novel environment-regulated genes in each cell type were enriched for genes that encode secreted factors, and cell-type-specific expression of their cognate receptors identified promising candidates for the modulation of learning and memory processes. Our characterization of the effect of novel experience on chromatin revealed thousands of cell-type-specific changes in chromatin accessibility. Coordinated analysis of chromatin accessibility and gene expression changes within individual cell types identified Fos/AP-1 as a key driver of novel experience-induced changes in chromatin accessibility and cell-type-specific gene expression. Together, these data provide a rich resource of hippocampal chromatin accessibility and gene expression profiles across diverse cell types in response to novel experience, a physiological stimulus that affects learning and memory.

## INTRODUCTION

The mammalian brain is comprised of neuronal and non-neuronal cell types that assemble into a network of functional neural circuits. In addition to cell-intrinsic developmental mechanisms, network assembly and refinement are driven in part by genetic programs that are activated in response to postnatal sensory experience.^1^ In this regard, post-mitotic neurons continue to adapt their gene expression programs to an ever-changing environment to encode novel experiences and, ultimately, memories. Both cell culture-based studies as well as *in vivo* stimulation paradigms, including light stimulation of the visual cortex of dark reared mice and seizure paradigms in the hippocampus, have shown that synaptic activity and/or growth factor signaling induces the transcription of a largely common set of early response genes (ERGs) that predominantly encode transcription factors (TFs) including FOS and EGR1.^2–6^ These ERG TFs in turn regulate a secondary wave of diverse cell-type-specific late-response gene (LRG) programs, which often encode synaptic effector proteins and secreted neuromodulatory proteins tailored to the function of specific cell types within neural circuits.^7–11^. A limitation of these prior studies is that the stimuli used to induce the gene expression programs have largely been non-physiological, and the specific features and diversity of gene expression programs that are activated by naturalistic stimuli remained to be identified.

Exposure to a novel naturalistic environment (NE) (also referred to as enriched environment) has gained attention as a mode of sensory stimulation that positively influences cognition, improves spatial memory and motor performance, and reduces anxiety.^12–14^ In rodents, continuous early-life NE exposure is associated with enhanced performance in learning and memory-associated tasks during adulthood.^14,15^ Moreover, prolonged novel environment exposure has also been shown to improve cognitive function during aging^16^ and both attenuate disease progression and ameliorate behavioral deficits in diverse mouse models of neurodevelopmental and neurodegenerative conditions.^17–19^ Previous efforts have characterized the gene expression outcomes that accompany structural changes after multiple weeks to months of NE exposure.^20–22^ However, the gene regulatory elements and the subsequent genes expression changes that initiate NE-dependent circuit remodeling remain largely uncharacterized. Indeed, to date, there has been little systematic exploration of acute gene expression programs in the hippocampus in response to naturalistic stimuli.

To bridge this gap in knowledge, we employed both region-specific RNA sequencing (RNA-seq) from the three main excitatory regions of the mouse hippocampus (CA1, CA3, and DG) and single-nucleus multiome sequencing (snMultiome-seq) to systematically characterize acute NE-responsive gene expression and chromatin accessibility patterns in the mouse hippocampus. We found that, whereas induction of ERG TFs was observed across all three hippocampal regions and in multiple cell types, novel experience-induced genes were expressed in a highly region- and cell-type-specific manner and were enriched for secreted proteins that function as neuromodulators and likely regulate neuronal connectivity and plasticity. We also profiled NE-responsive genomic cis-regulatory regions, uncovering a widespread role for AP-1 factors in these cell-type-specific responses. Taken together, our findings identify the chromatin accessibility and gene expression landscape following NE exposure across 14 cell types of the hippocampus and provide a rich resource for the functional interrogation of activity-dependent genes and the gene regulatory elements that drive their expression in the context of spatial learning and memory paradigms. These datasets are accessible via our accompanying web-based searchable database, where users can input gene names and extract plots of gene expression and AP-1 accessibility across all cell classes and times of NE exposure (https://greenberg.hms.harvard.edu/project/novel-environment-gene-database/).

## RESULTS

### Transcriptional landscape following NE exposure

To characterize hippocampal gene expression kinetics in response to brief NE exposure, we transferred adult C57BL/6J mice from their home cage (HC) to a novel context for 30 min, followed by a return to HC (**Fig. 1A**). Immunostaining for FOS and EGR1, two well-characterized ERG TFs, confirmed robust ERG induction within two h of NE exposure (**Fig. 1B**, **Sig. S1A**), as expected.^6,20,21,23–25^ Having confirmed the efficiency of this stimulation paradigm, we employed two complementary approaches to systematically characterize NE-induced gene expression programs. Bulk RNA-seq was used to achieve deep sequencing coverage of both nuclear and cytoplasmic RNA from samples collected following a time course of brief NE exposure. To this end, mice were sacrificed at ten time points, spanning a 24 h period following 30 min NE exposure. Hippocampal tissue was micro-dissected into CA1, CA3, and dentate gyrus (DG) regions and libraries prepared for deep sequencing (**Fig. S1B**). Principal component analysis (PCA) effectively separated bulk RNA-seq samples by anatomical region, highlighting cell-class-specific gene expression programs and the reproducibility of our microdissection (**Fig. 1C**). To profile cell-type-specific gene expression and chromatin accessibility changes, hippocampal nuclei were isolated 1 h post-NE exposure for single-nucleus 10X multiome (snRNA + snATAC) sequencing. Following dimensionality reduction, the BRAIN Initiative Cell Census Network (BICCN) snRNA-seq atlas was used to assign 14 major cell populations (**Fig. 1D**), which could be further clustered into 33 sub-class cell populations (**Fig. S2A**). After applying quality control metrics (see Methods, **Fig. S2B-D**), we obtained transcriptomes from a total of 45,844 cells, with the resulting cell clusters well integrated between HC and NE samples (**Fig. S2B**).

**Figure 1:**
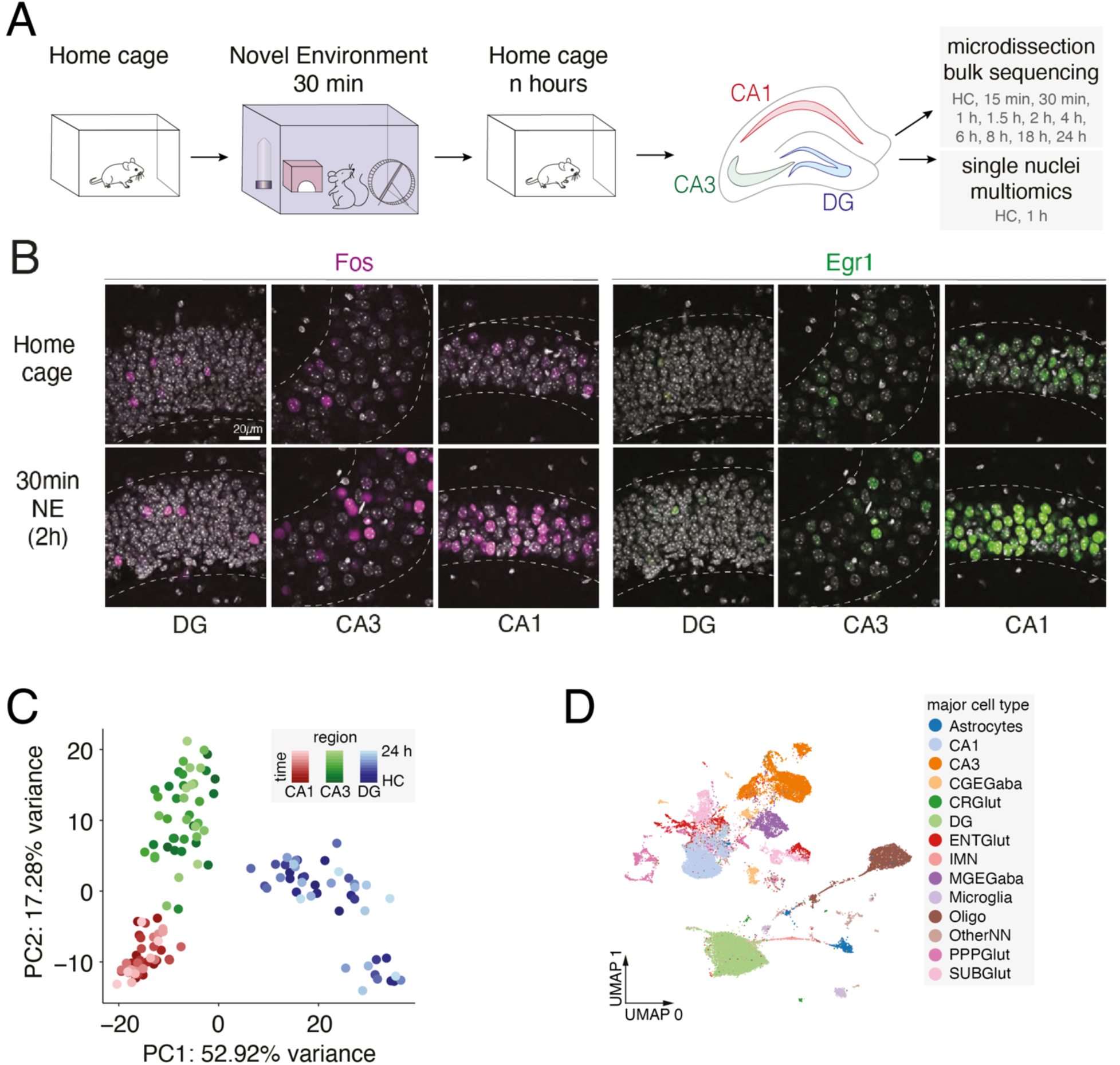
NE exposure stimulates hippocampal ERG activity. **(A)** Overview of experimental design. **(B)** Representative immunofluorescence images of FOS and EGR1 protein levels in CA1, CA3, and DG from HC and 90 min following a 30 min NE exposure. Scale bar 20 µm **(C)** Principal component (PC) analysis of read depth-normalized RNA-seq counts from the top 2000 most variable genes across CA1, CA3, and DG. Each point is a sample colored by region and time following a 30 min NE exposure. HC = home cage. **(D)** UMAP visualization of snRNA-seq from nuclei from all collected timepoints with cell type information overlaid. CGEGaba: GABA-ergic neurons originating from the caudal ganglionic eminence (e.g. VIP and CCK), CRGlut: Cajal-Retzius glutamatergic neurons, ENTGlut: entorhinal glutamatergic neurons, IMN: immature neurons (e.g. adult-born stem cells), MGEGaba: Gaba-ergic neurons originating from the medial ganglionic eminence (e.g. PV and SST), Oligo: oligodendrocytes, OtherNN: other non-neuronal cells, PPPGlut: glutamatergic neurons originating from the presubiculum, parasubiculum, and postsubiculum, SUBGlut: glutamatergic neurons originating from the subiculum. n = 45,844 cells, 3 mice.

### Region-specific differences in home cage gene expression

We further characterized region-specific gene expression differences in HC animals as an additional test of data quality. Hierarchical clustering of differentially expressed genes identified in pairwise comparisons between HC CA1, CA3, and DG bulk RNA-seq samples was used to define region-enriched genes (**Fig. 2A**, **Fig. S3A**). Our snRNA-seq dataset confirmed that excitatory neurons were major contributors to region-specific gene expression differences. In addition, within CA1 and CA3, MGE-derived GABAergic neurons (e.g. PV and SST inhibitory neurons) contribute to the region-specific differences, whereas expression signatures characteristic of immature neurons were most highly detected in DG, an area known for containing a stem cell niche for the generation of adult-born neurons (**Fig. 2A**). Overall, these cell-type- and region-specific gene expression differences strongly align with the results of prior studies (**Fig S3B**),^26,27^ with CA1-, CA3-, and DG-enriched gene expression signatures reflecting their specialized roles in synaptic signaling and neuronal function (**Fig. 2B & C, S3C**) that are then further modified by neuronal activity.

**Figure 2:**
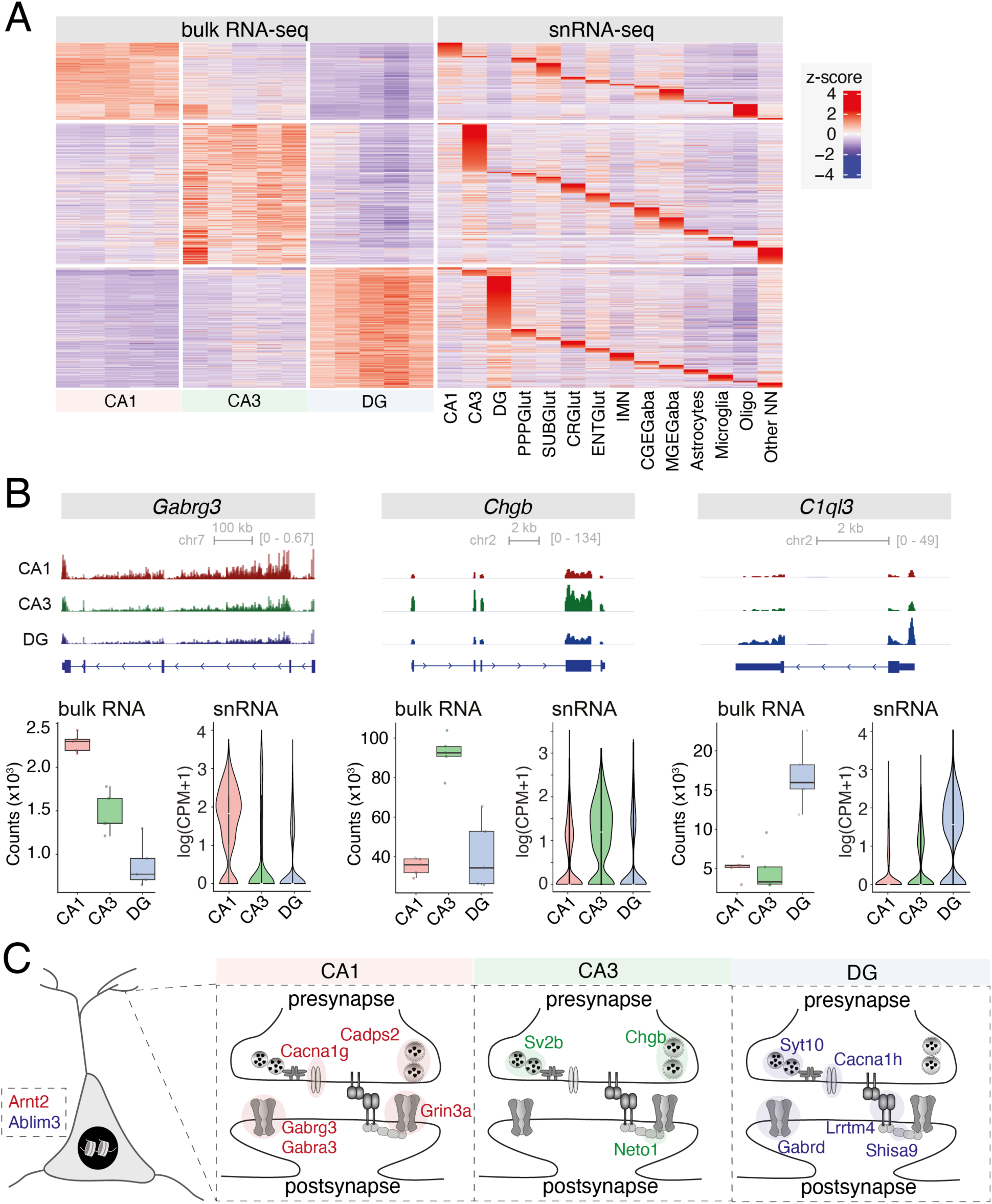
Region-specific differences in HC gene expression. **(A)** Heatmap of row-normalized z-scores from bulk RNA-seq and snRNA-seq expression across all regions and major cell types, respectively. Each bulk RNA-seq column is a biological replicate (n=5) separated by region. Bulk RNA-seq CA1-, CA3-, and DG-enriched genes were determined by k-means clustering. **(B)** Merged genome browser tracks (top), box and whisker plots (bottom left) of bulk RNA-seq depth-normalized counts, and violin plots (bottom right) of snRNA-seq counts for *Gabrg3* (CA1- enriched), *Chgb* (CA3-enriched), and *C1ql3* (DG-enriched). Box plots are shown as median ± IQR (whiskers = 1.5*IQR). Violin plots are shown with mean as a white dash. **(C)** Schematic of region-enriched synaptic genes in CA1, CA3, and DG.

### Time-resolved NE-driven gene expression changes

Having characterized basal hippocampal region-specific gene expression differences, we next investigated the nature of NE-driven gene expression responses. Differential gene expression analysis comparing each region-specific bulk RNA-seq dataset to the appropriate HC sample revealed 704 significant differentially expressed genes (**Fig. 3A, S4A, Table 1**, P_adj_ < 0.05 and abs (fold change) > 1.2, see Methods), whereas snRNA-seq analysis after 1h of NE exposure identified 3,458 genes to be differentially regulated (**Fig. 3B**, **Table 1**, P_adj_ < 0.05 and abs (fold change) > 1.2). Of the 704 differentially regulated genes in our bulk sequencing comparisons, 570 genes show significant differential expression within 2 h of stimulus onset (**Fig. 3C, S4B**), with the early-response genes being enriched for GO terms related to transcription factor activity (**Fig. 3D, S4C, Table 2**). Clustering these genes based on their region-specific induction kinetics revealed a subset of early-response genes that were significantly induced across all three regions (**Fig. 3C, D**), including well-characterized ERG TF family members (e.g. *Fos, Fosb, Npas4, Nr4a1*, and *Egr3*) (**Fig. 3D, E**). By contrast, other early-response TFs exhibited regional specificity, including the ERG TFs *Nr4a3*, *Atf4*, and *Klf4* (CA1-induced), the bHLH TF *Atoh8* (CA3-induced), and *Bcl6* (DG-induced), a transcriptional repressor of multiple developmental pathways that is thought to promote neurogenesis^28^ (**Fig. 3D, F**). We also identified many NE-induced synaptic factors, including the plasticity-related proteins *Arc* and *Homer1* (common), the synaptic vesicle protein *Synpr* (CA1-induced), the regulator of synaptic membrane exocytosis *Rims4* (CA3-induced), and *Dact1* (DG-induced), a member of the Wnt pathway (**Fig. S4B**). Activity-regulated genes that showed significant expression changes in both bulk and scRNA-seq were mostly regulated in excitatory neurons, with a subset showing induction across inhibitory and non-neuronal cell types (**Fig. S4D**).

**Figure 3:**
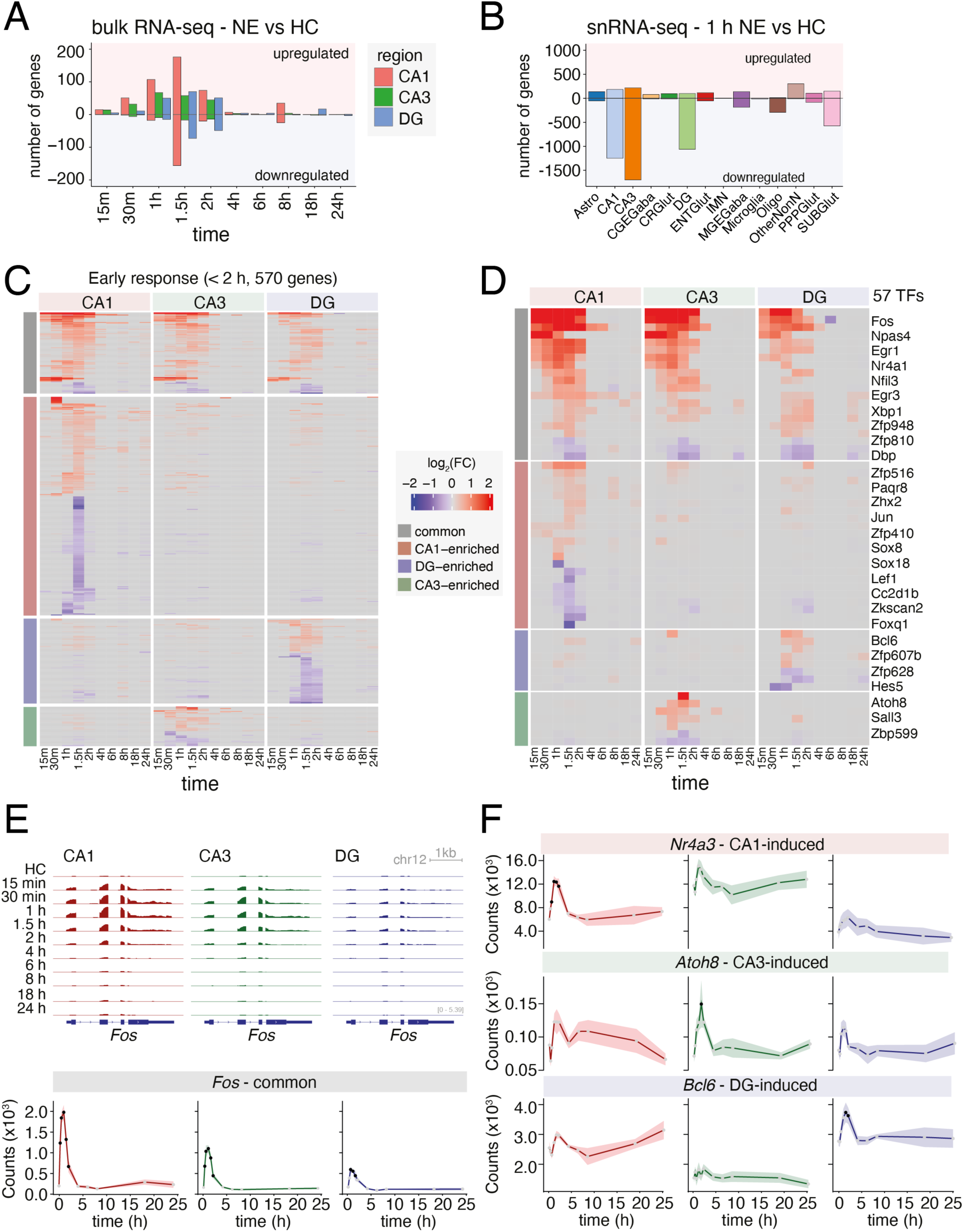
NE-driven gene expression changes. **(A)** Bar plot of significant (DESeq2 P_adj_ < 0.05, |FC| > 1.2) differentially expressed genes in CA1, CA3, and DG by bulk RNA-seq following a time course of NE exposure. Upregulated and downregulated genes are plotted as positive and negative values on the y-axis, respectively. **(B)** Bar plot of significant (Wilcoxon Rank Sum Test P_adj_ < 0.05, |FC| > 1.2) differentially expressed genes in all major cell types by snRNA-seq following 1 h brief NE exposure (30 min NE, followed by 30 min back in HC). Upregulated and downregulated genes are plotted as positive and negative values on the y-axis, respectively. **(C)** Heatmap of log_2_(fold change) in gene expression relative to HC at each time point of NE exposure in CA1, CA3, and DG by bulk RNA-seq. Genes were defined as early (< 2 h)-response genes based on the first time point at which |FC| > 1.2 and separated into CA1- CA3- and DG-enriched or Common based on whether fold change requirements were satisfied in one or more than one region, respectively. (**D**) Heatmap as in (C) for early-response genes that are annotated Transcription Factors (TFs). For clarity, every other row has been labeled. A full list of genes and their regulatory category can be found in Table 1. **(E)** Merged genome browser tracks (top) and line plot of bulk RNA-seq depth-normalized counts for the ERG TF *Fos* following NE exposure in CA1, CA3, and DG. In line plot, dark line = mean counts, shading = ±SEM, black point = DEseq2 P_adj_ < 0.05, light gray point = DEseq2 P_adj_ > 0.05. **(F)** Line plots of bulk RNA-seq depth- normalized counts for *Nr4a3* (CA1-induced), *Atoh8* (CA3-induced), and *Bcl6* (DG-induced) following NE exposure in CA1, CA3, and DG. In line plots, dark line = mean counts, shading = ±SEM, black point = DEseq2 P_adj_ < 0.05, light gray point = DEseq2 P_adj_ > 0.05.

Coordinated ERG expression has been observed in individual cells in the visual cortex in response to light stimulation following dark rearing.^5^ We therefore analyzed the correlation of commonly induced ERG expression across single excitatory neurons in CA1, CA3, and DG. ERGs exhibited significantly stronger pair-wise correlations within individual cells compared to similarly expressed, non-induced genes (**Fig. S4E**). Our results thus show that coordinate ERG expression is a consistent phenomenon across multiple brain regions and occurs in response to a subtle, naturalistic stimulus such as exposure to a novel environment.

ERG TFs induce a second wave of LRGs that drive structural and functional changes related to plasticity.^29,30^ We identified 134 LRGs by bulk RNA-seq that showed significant differential expression ≥ 2 h after brief NE exposure (**Fig. 4A**, P_adj_ < 0.05 and abs (fold change) > 1.2, see Methods). Most LRGs are specifically induced in one hippocampal region, such as *Dclk3* (DG-induced), a serine-threonine kinase thought to regulate the activity of many TFs and chromatin remodelers,^31^ *Mgp* (CA3-induced), a matrix-associated protein and regulator of the BMP growth pathway,^32^ and *Ttl* (CA1-induced), which enables tubulin-tyrosine ligase activity that is required for microtubule organization and axon extension (**Fig. 4B**).^33^ Overall, upregulated LRGs were enriched for functions related to extracellular secretion (**Fig. S4B, Table 2**), such as *Ccn1* (common), a secreted factor involved in cell adhesion, migration, and neuroinflammation processes critical for synaptic plasticity and tissue repair, as well as *Inhba* (CA1-induced), a member of the transforming growth factor-β (TGFβ) superfamily, *Plat* (CA3-enriched), which encodes the protein tPA that cleaves neurotransmitter receptors, and the endoplasmic reticulum calcium-binding chaperone *Calr* (DG-induced) (**Fig. 4C**). Novel environment exposure thus triggers region-specific LRG programs that may promote circuit-specific synaptic plasticity.

**Figure 4:**
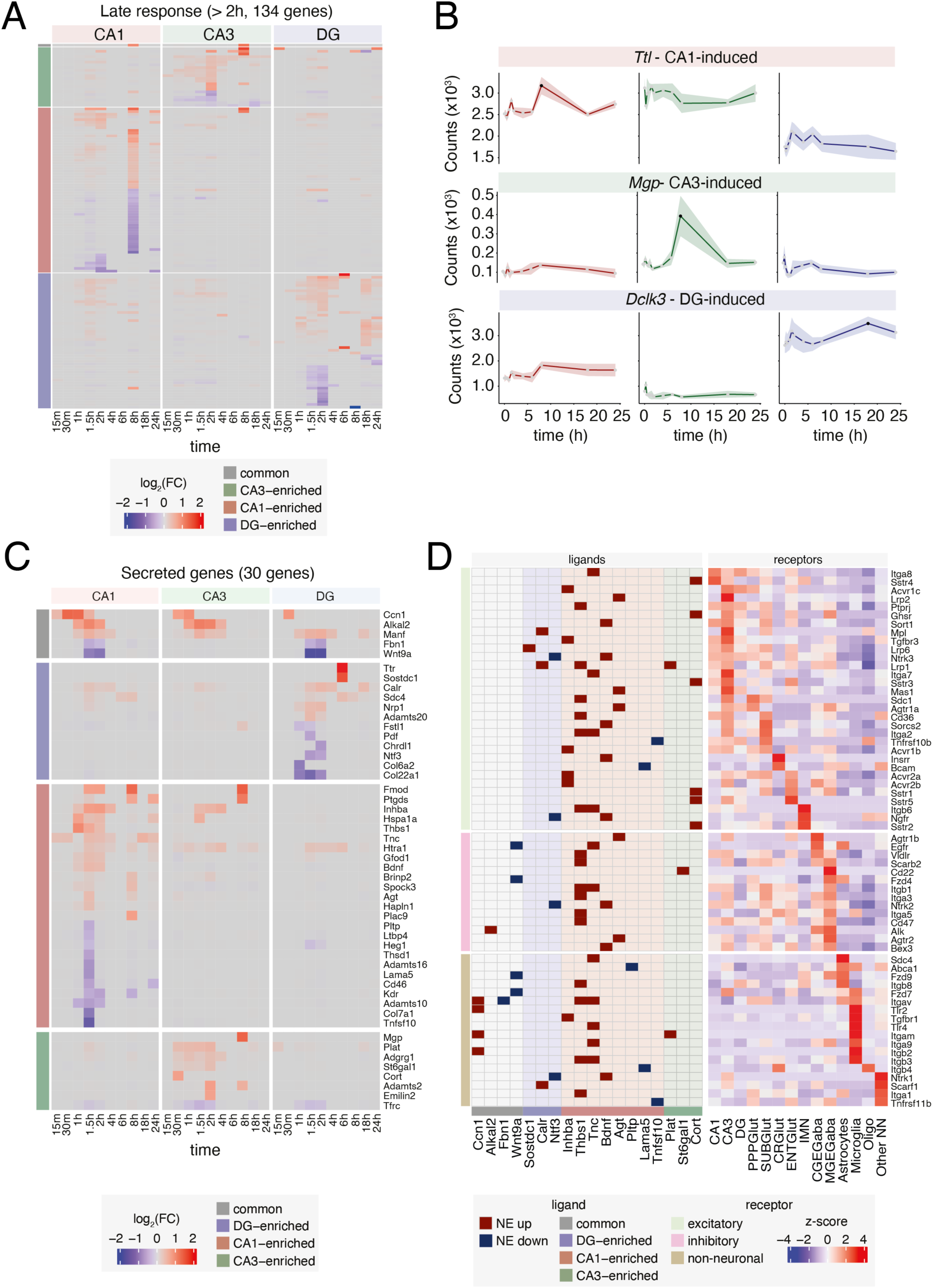
NE-driven late-response genes. **(A)** Heatmap of log_2_(fold change) in gene expression relative to HC at each time point of NE exposure in CA1, CA3, and DG by bulk RNA-seq. Genes were defined as late (> 2 h)-response genes based on the first time point at which |FC| > 1.2 (excluding early-response genes from Figure 3C) and separated into CA1- CA3- and DG-enriched or Common based on whether fold change requirements were satisfied in one or more than one region, respectively. (**B**) Line plots (top) and merged genome browser tracks (bottom) of bulk RNA-seq depth-normalized counts for *Ttl1, Mgp,* and *Dclk3* following NE exposure in CA1, CA3, and DG, respectively. In line plot, dark line = mean counts, shading = ±SEM, black point = DEseq2 P_adj_ < 0.05, light gray point = DEseq2 P_adj_ > 0.05. **(C)** Heatmap of log_2_(fold change) in gene expression of all significant NE-regulated genes that are annotated secreted factors relative to HC at each time point of NE exposure in CA1, CA3, and DG by bulk RNA-seq. Colors over gene names indicate that the gene has at least one known receptor. **(D)** Heatmap of secreted factors (ligands) from (C) with their receptors. Left grid shows ligand-receptor pair, with dark shading indicating whether ligand expression was upregulated (dark red) or downregulated (dark blue) in response to NE. Background shading indicates the hippocampal region of ligand NE gene expression regulation (common, CA1- CA3- or DG-enriched). Right heatmap shows row- normalized z-scores of cognate receptor expression from HC snRNA-seq across all major cell types, ordered by cell type of maximum receptor expression.

To explore whether secreted factors induced in excitatory neurons could influence neighboring neuronal and non-neuronal cell types, we mapped the expression of known cognate receptors^34^ of NE-dependent secreted factors across cell types in our single-nucleus RNA-seq (snRNA-seq) data, revealing receptor expression across multiple hippocampal cell types (**Fig. 4D**). Interestingly, the receptors for *Ccn1*, a NE-induced cell adhesion factor, are mainly expressed in microglia, consistent with prior reports that CCN1 activates this cell type.^35^ This tight temporal regulation of *Ccn1* expression may contribute to proper hippocampal function insofar as temporally constrained microglial activation is important for dendritic spine phagocytosis, whereas prolonged activation of microglia has been associated with neurodegenerative conditions.^36^ By contrast, brain-derived neurotrophic factor (*Bdnf,* CA1-upregulated) and neurotrophin-3 *(Ntf3*, DG-downregulated) bind to receptors in excitatory, inhibitory, and non-neuronal cells, which may in part explain their widespread roles in promoting neuronal survival, differentiation, and synaptic connectivity.^37,38^ Intriguingly, we find that receptors for CA3-induced ligands tend to be restricted to excitatory neurons, whereas receptors for CA1-induced ligands (e.g., *Bdnf* and *Inhba*) are broadly expressed across many cell types, suggesting that secreted factors that are novel experience-induced in excitatory neurons may coordinate a wide range of cellular responses to promote aspects of learning and memory, not only in neighboring neurons but also non-neuronal cells.

### Cell-type-specific NE-induced gene expression programs

Further cell type resolution was provided by our snRNA-seq data, with genes classified based on whether their expression changes were common across multiple cell classes (significant in any combination of two cell types: excitatory, inhibitory, or non-neuronal), or specifically enriched in one of these classes. Although we identified substantially more downregulated genes compared to upregulated genes in many cell types, NE-induced genes showed higher fold changes compared to downregulated genes (**Fig. S5A**). We observed stimulus-dependent gene expression changes across excitatory, inhibitory, and non-neuronal cell types: 269 genes showed common regulation across multiple cell classes, 1,042 genes were excitatory-enriched, 68 genes were inhibitory-enriched, 366 genes were enriched in non-neuronal cells, and 26 genes were enriched in both inhibitory and non-neuronal cells (**Fig. 5**). For example, Protocadherin 7 (*Pcdh7*), a cell adhesion molecule required for axonal and dendritic outgrowth and function was induced in MGE-derived parvalbumin+ and somatostatin+ inhibitory neurons.^39^ By contrast, astrotractin2 (*Astn2*), which is required for neuronal migration and is a regulator of synaptic strength and whose loss of function contributes to symptom onset of neurological disorders was induced in CGE-derived vasoactive intestinal polypeptide (VIP)+ and cholecystokinin (CCK)+ interneurons.^40,41^ The voltage-dependent L-type calcium channel subunit alpha-1C (*Cacna1c*), which mediates calcium ion influx and the reactivation of astrocytes, was upregulated in astrocytes.^42^ Finally, *Adipor2* (Adiponectin receptor protein 2), a key hormone for the regulation of glucose and lipid metabolism^43^ that has been linked to reduced extinction learning in contextual fear conditioning is selectively induced in oligodendrocytes.^44^ Our findings indicate that different cell classes in the hippocampus have specialized functions in responding to NE exposure, with excitatory neurons showing the most pronounced induction of gene expression and non-neuronal cells and inhibitory neurons displaying significant responses as well.

**Figure 5:**
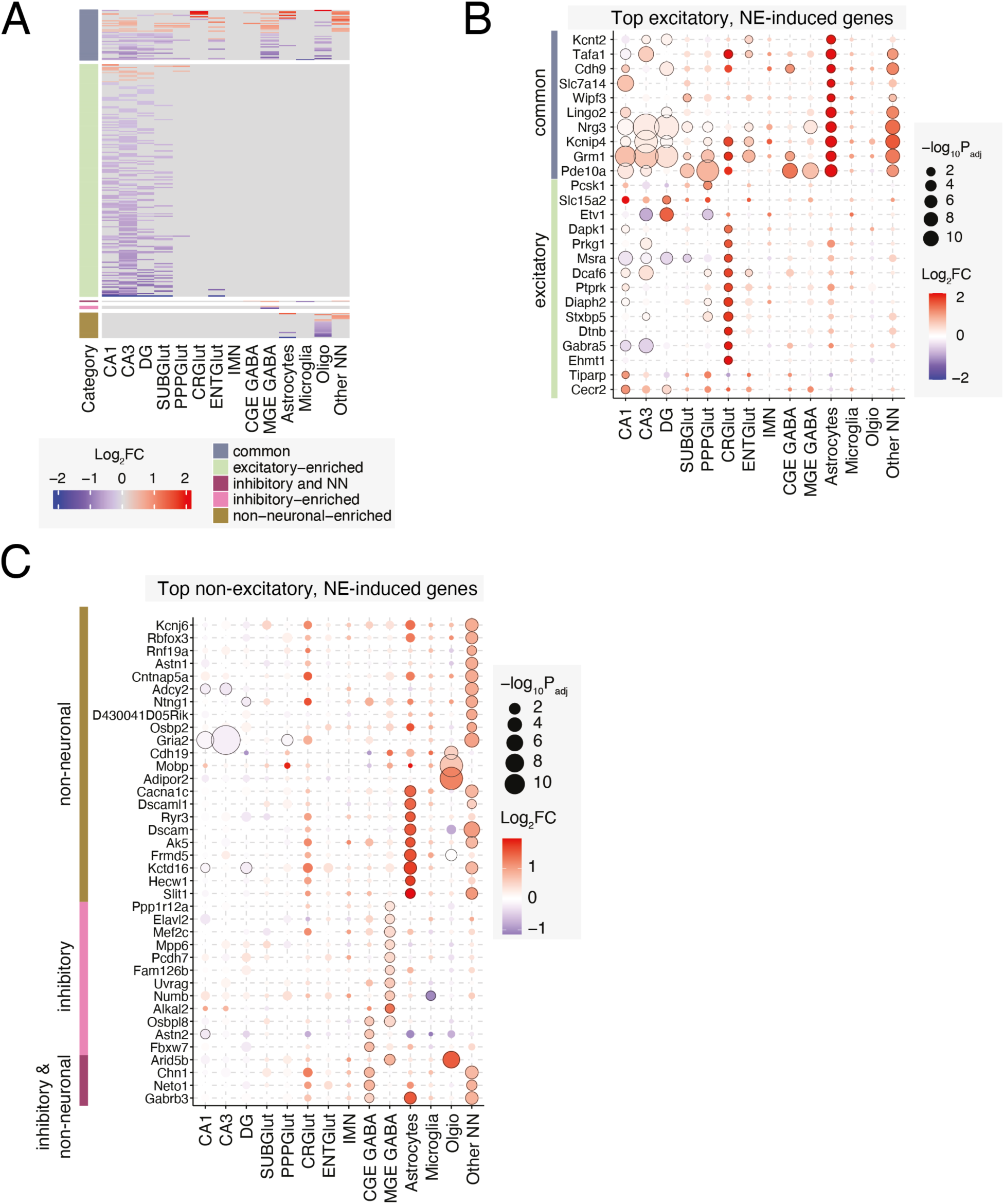
Cell-type-specific NE-induced gene expression programs. **(A)** Heatmap of log_2_(fold change) in gene expression relative to HC at 1 h following brief NE exposure in all major cell types by snRNA-seq. Genes were separated based on their induction in one or more cell classes (excitatory, inhibitory, or non-neuronal). **(B)** Dot plot of the top common or excitatory-specific induced genes (color scheme as in (A). Dot color indicates log_2_(fold change), size indicates - log_10_(P_adj_). Black outline indicates P_adj_ < 0.05. (**C**) Dot plot of the top induced genes in each inhibitory or non-neuronal cell type that are not significantly induced in any excitatory cell type. Dot color indicates log_2_(fold change), size indicates -log_10_(P_adj_). Black outline indicates P_adj_ < 0.05.

### Continuous NE exposure boosts expression of LRGs

We were surprised to find that very few previously characterized LRGs showed significant gene expression changes 6 h after a brief (30 min) exposure to NE (**Fig. 3A**), as robust LRG expression at this time point has previously been observed in cultured neurons exposed to membrane depolarizing agents^8,9^ and growth factor stimulation^45,46^ as well as *in vivo* in response a kainic acid-induced seizure.^6,25^ To determine if exposure to a novel environment for a longer period of time was more effective at inducing LRGs in the hippocampus, we transferred adult C57BL/6J mice from their home cage (HC) to a novel context for 6 h (cNE) and prepared bulk or snMultiome-seq libraries as described above (**Fig. 6A, S5B**). This longer exposure to novelty led to the identification of differentially expressed genes in both bulk RNA-seq and snRNA-seq (2,371 and 1,275 genes, respectively) (**Fig. 6B, S5C**). As observed in response to brief NE, LRGs in response to cNE were largely non-overlapping between CA1, CA3 and DG (**Fig. 6C, D**). Bulk RNA-seq showed more LRGs induced in the DG compared to CA1 and CA3 in response to cNE (**Fig. 3A, 6B**). Upregulated genes were enriched for GO terms associated with responses to extracellular stimuli, while downregulated genes were enriched for terms related to synaptic function and energy metabolism, particularly in the DG (**Fig. 6E**). Intriguingly, we identified secreted factors whose expression was modulated by cNE as measured by bulk RNA-seq, which revealed both overlapping (*Inhba, Wnt9a*) and distinct (*Fbn1, Lama1, Slit2*) ligands compared to brief NE (**Fig. 6A, S6A-C)**. These ligands bind receptors that are expressed across diverse excitatory, inhibitory and non-neuronal cell types, suggesting their potential to regulate tissue-wide cellular function in response to NE (**Fig. S6A, B**).

**Figure 6:**
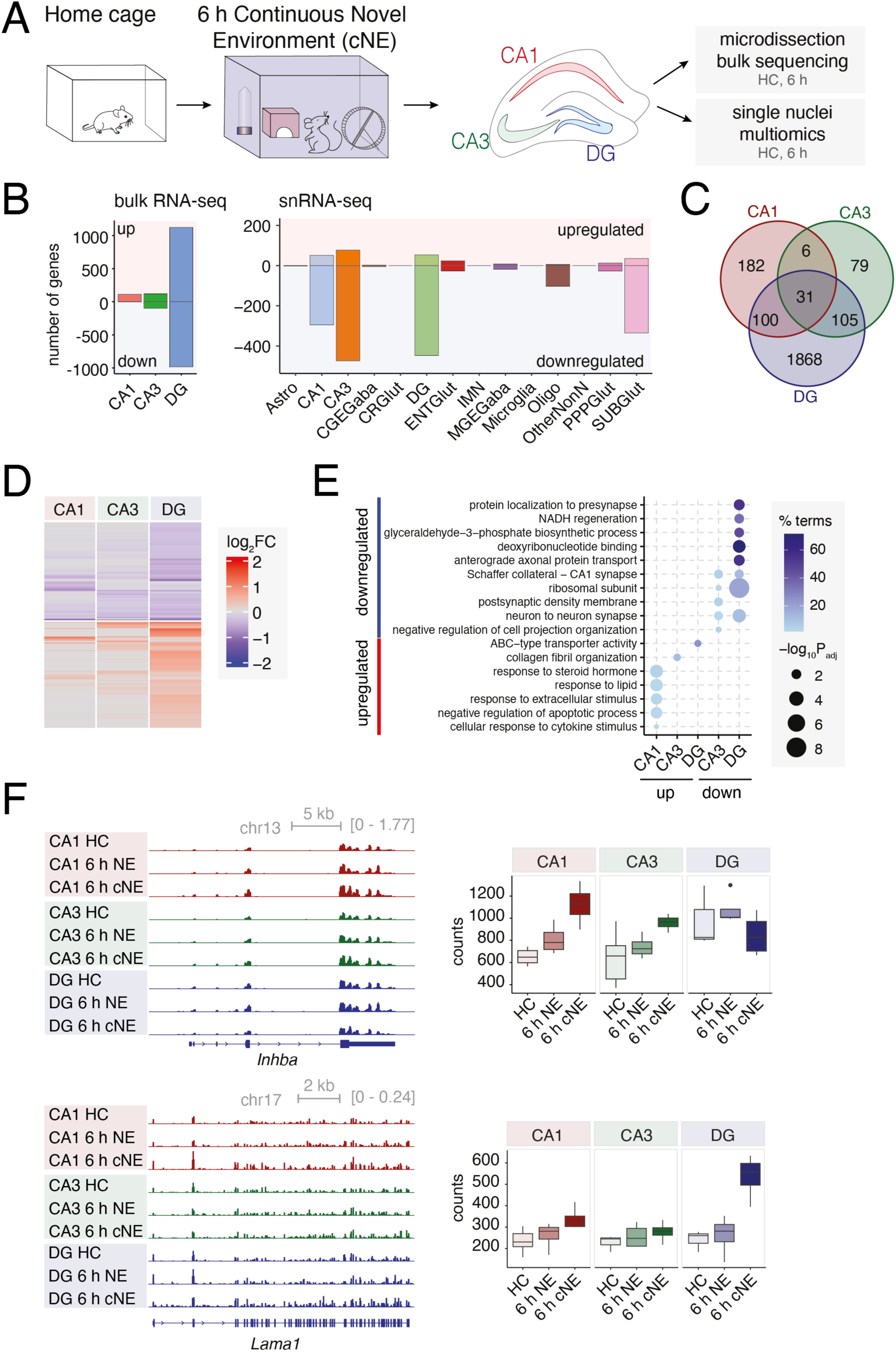
Continuous NE exposure boosts expression of LRGs. **(A)** Overview of experimental design. **(B)** Left: bar plot of significant (DESeq2 P_adj_ < 0.05 & |FC| > 1.2) differentially expressed genes in CA1, CA3, and DG following cNE exposure. Right: bar plot of significant (Wilcoxon Rank Sum Test P_adj_ < 0.05, |FC| > 1.2) differentially expressed genes in all major cell types by snRNA-seq following cNE exposure. Upregulated genes and downregulated genes are plotted as positive and negative values on the y-axis, respectively. **(C)** Venn diagram of overlap in DE genes following cNE exposure in CA1, CA3, and DG. **(D)** Heatmap of log_2_(fold change) in gene expression relative to HC in response to cNE exposure in CA1, CA3, and DG. Genes were separated into upregulated or downregulated clusters by k-means hierarchical clustering. **(E)** Dot plot of the top enriched GO terms for upregulated and downregulated genes in (D), separated by hippocampal region. **(F)** Merged genome browser tracks (left), box and whisker plots of bulk RNA-seq depth-normalized counts (right) for *Inhba* and *Lama1* following brief 6h and continuous 6h novel environment exposure (bNE and cNE, respectively). Box plots shown as median ± IQR (whiskers = 1.5*IQR).

We considered whether the differences between brief and continuous NE reflected fundamentally distinct gene expression programs induced by different stimulation durations,^8^ or whether continuous NE leads to a cumulative increase in gene expression over time as RNA gradually accumulates in the cytoplasm. In this regard, we found that the directionality of expression changes between brief NE and cNE were not markedly different. Rather, the magnitude of expression changes was greater in response to cNE. For example, *Inhba,* a gene involved in hormone secretion, was significantly induced in CA1 in response to both brief and continuous NE, but we observed a larger induction following 6 h of cNE (**Fig. 6F**). In contrast, we observed a significant induction of *Lama1* in DG, but only in response to cNE, where this gene encodes a laminin subunit thought to mediate cell attachment and migration.^47^ Moreover, focusing on our snRNA-seq datasets, which preferentially capture newly transcribed RNA, we found largely overlapping gene expression changes between brief and continuous NE exposure (**Fig. S6D**), consistent with similar gene expression responses to these two conditions. The strong correlation between nuclear RNA and total RNA changes observed at 1h was no longer observed after 6 h of continuous NE exposure (**Fig. S5D**), further arguing that the gene expression changes observed by bulk RNA-seq are due to a shift in cytoplasmic mRNA levels.

Together, these data support the hypothesis that cNE causes repeated activation of activity-dependent ERGs, leading to a gradual cytoplasmic build-up of late-response transcripts that may be important for shaping long-term hippocampal circuit connectivity. Notably, we further compared our cNE bulk dataset to gene expression changes in response to kainic acid-induced seizures, a commonly used stimulus in genomic identification of activity-dependent gene expression.^6^ These stimulation paradigms induce vastly different magnitudes and directionality of gene regulation. When we focused our attention on a select list of kainic acid-induced candidate genes, we discovered that NE also induced genes such as *Scg2, Inhba, Pcsk1 or Nptx2* but downregulated *Lonrf1, Cgref1* and *Adpgk* (**Fig. S7**). This differential response suggests that, while seizures trigger a broad and robust activation of genes linked to stress and excitotoxicity, novel environment exposure may instead promote more selective transcriptional programs, potentially favoring pathways involved in neuroplasticity, adaptation, and homeostasis.

### Accessibility of AP-1 sites predicts NE-induced gene upregulation

We next examined NE-regulated chromatin accessibility changes in our snMultiome datasets to gain insight into the regulatory mechanisms that mediate these novel experience-activated gene programs. In this regard, cell clustering based on chromatin accessibility largely matched that based on gene expression (**Fig. S8A**), where cell clusters from distinct NE time points were well integrated and passed all quality control metrics (**Fig. S8B**). Differential accessibility analysis identified thousands of regions exhibiting altered accessibility in response to 1 h brief NE and 6 h cNE relative to HC across different cell types, with many NE-regulated peaks showing strong specificity for only one or a few cell types (**Fig. 7A**). For example, ATAC peaks surrounding the *Scg2* gene body showed a mix of three types of peaks: 1) putative enhancer regions that are constitutively open across many cell types 2) cell class-specific, constitutively open regions 3) NE-induced open regions that are selective for CA1 pyramidal neurons (**Fig. 7B**).

**Figure 7:**
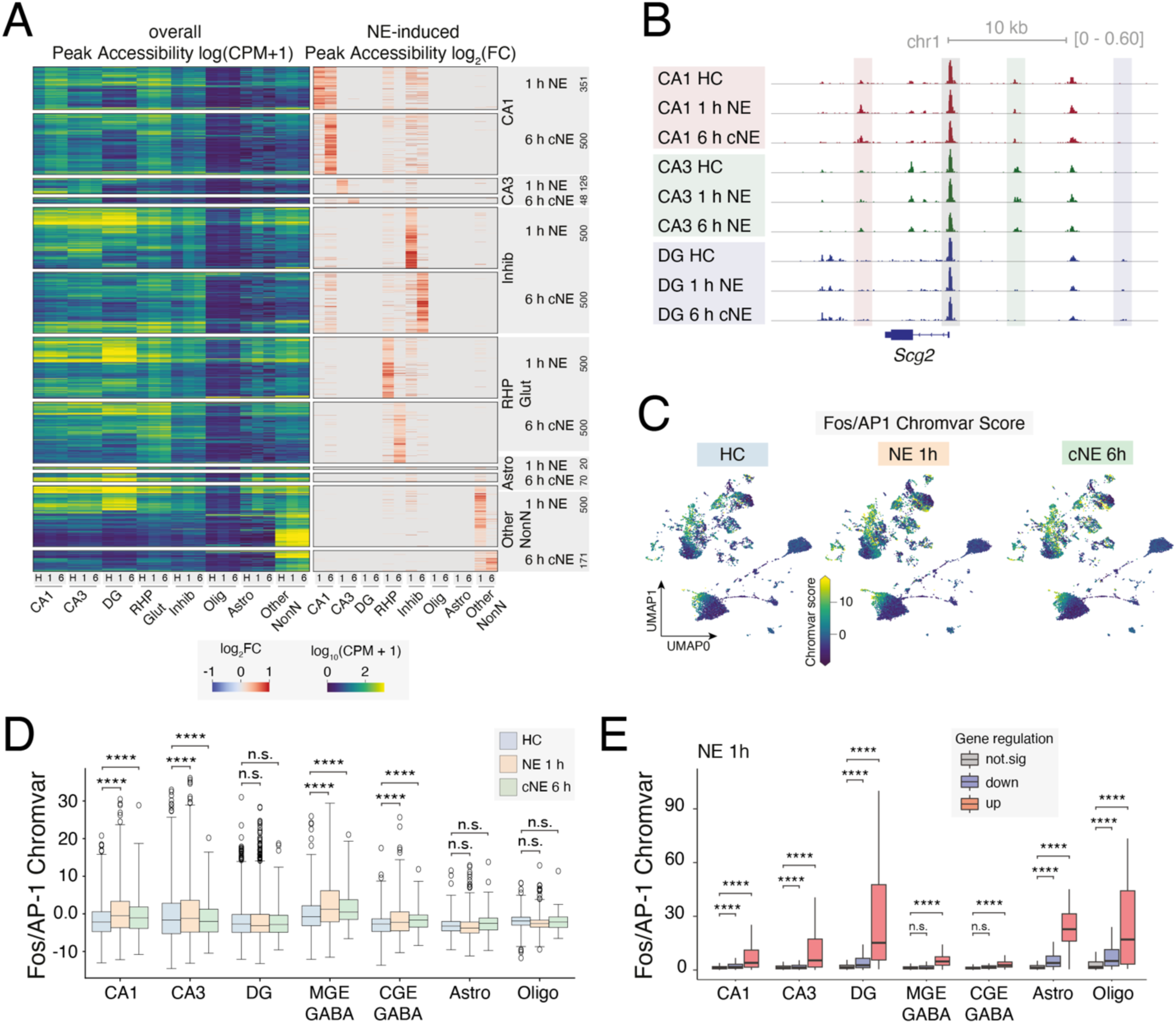
Chromatin accessibility changes in response to NE at single-cell resolution. **(A)** Heatmap of overall log_10_(counts per million) peak accessibility (left) and NE-induced log_2_(fold change) (right) in HC, 1 h brief or 6 h continuous NE stimulation (H, 1, & 6, respective) of all major cell types in snATAC-seq. Clusters were defined using the BICCN Brain Atlas. CA1 & CA3 = excitatory neurons in CA1 and CA3, respectively. Inhib = inhibitory neurons. RHP Glut = retrohippocampal glutamatergic neurons. Astro = astrocytes. Other NonN = other non-neuronal cell types. Numbers next to labels indicate number of peaks in each category. (**B**) Genome browser tracks of chromatin accessibility merged by cell type surrounding the *Scg2* locus. Red, green and blue shading indicates a region of cell-type-specific chromatin opening in HC and in response to brief 1 h and continuous 6 h (cNE) novel environment exposure in CA1, CA3 and DG, respectively. Grey shading indicates *Scg2* promoter accessibility. **(C)** UMAP visualization of Fos/AP-1 Chromvar score (see Methods). **(D)** Box and whisker plot of Fos/AP-1 Chromvar scores within select major cell types. **(E)** Fos/AP-1 Chromvar score separated by up- (red), down- (blue), or not significant (grey) regulation of gene expression changes in sn-RNAseq after 1 h NE in select cell classes. **** = P < 0.05, Mann Whitney U test followed by BH multiple hypothesis correction

Motif enrichment analysis revealed strong enrichment for a number of ERG TF-binding motifs at sites that exhibited increased accessibility in CA1 and CA3 pyramidal neurons in response to NE exposure (**Fig. S9A**). Notably, however, we did not observe enrichment of these motifs in NE-exposed samples within DG neurons, consistent with the lower inducibility of NE-responsive gene programs in this region. Strikingly, the AP-1 motif exhibited the highest enrichment in all neuronal cell types, including inhibitory cells, with excitatory neurons, particularly those in CA1, showing the largest increase in AP-1 motif accessibility in response to NE (**Fig. 7C, D, S9B, S10A**). To investigate whether an increase in AP-1 motif accessibility was predictive of gene induction, we first binned cells into quantiles based on the Chromvar scores,^48^ where a higher score indicates greater NE-induced accessibility of AP-1 motifs in that cell (**Fig. S10B,** single gene example; Table 3).

We then examined NE-induced gene expression changes in cells with high versus low Chromvar score across all cell classes. Strikingly, only AP-1 target genes that showed robust induction by bulk RNA-seq in response to cNE exposure (**Fig. S7**) also showed strong AP-1 Chromvar activation scores, whereas non-regulated or downregulated genes did not (**Fig. S10C).** More broadly, as analyzed by snRNA-seq, genes that were upregulated following NE exposure had overall higher AP-1 Chromvar scores in their respective cell types (**Fig. 7E, S10D**). These findings indicate a robust relationship between NE-induced opening of AP-1 motifs and regulated gene expression, reinforcing the idea that AP-1 motif accessibility is a strong predictor of NE-induced gene expression.

Overall, our data suggest that the presence of accessible AP-1 motifs is a critical factor for determining the transcriptional response to NE in many cell types and that Fos binding to these sites likely mediates gene induction by these regulatory elements across cell types. More generally, our findings reveal three distinct classes of chromatin accessibility: regions that are universally open across all cell types, those specific to certain cell types under baseline conditions, and regions that become accessible only in response to novel environments, often in a cell-type-specific manner. The diversity of these accessible chromatin regions in our dataset further highlights the potential of gene regulatory elements to undergo chromatin remodeling in response to not only basal neuronal circuit activity, but also to acute physiological stimuli.

## Discussion

Exposure to novel environments modulates neural connectivity over time, leading to improved cognitive function or attenuation of neurological disease progression. This structural and functional plasticity involves key regions of the hippocampus, including DG, CA3, and CA1, which have distinct but interconnected roles in encoding, processing, and storing information. Previous studies have demonstrated that ERG induction occurs in these regions in response to NE exposure, yet the downstream transcriptomic changes have largely been examined only after days to weeks of prolonged exposure, leaving a gap in our understanding of the molecular mechanisms by which NE by activating ERGs acutely regulates neuroplasticity. ^20,22^

Using a combination of bulk RNA-seq and single-nucleus multiomic sequencing (snRNA-seq + snATAC-seq), our investigation fills a critical gap in knowledge by focusing on the gene regulatory and mRNA transcript changes that occur in mouse DG, CA3, and CA1 during the first 24 h after NE exposure. We observe a rapid induction of ERG TFs across all three hippocampal regions and multiple cell types within 2 h of NE exposure. In contrast, LRGs exhibited region- and cell-type-specific expression patterns, including genes enriched for protein functions related to secreted factors that signal to both neuronal and non-neuronal cells. Prolonged NE exposure led to a more pronounced induction of LRGs when measuring both nuclear and cytoplasmic RNA levels by bulk RNA-seq, but we did not observe a corresponding increase in transcription of these genes as measured by snRNA-seq. This suggests that sustained NE exposure drives repeated activation of these transcriptional programs leading to the cytoplasmic accumulation of LRG transcripts.

Our multiomic data also revealed that novel environment (NE) exposure induces cell type-specific changes in chromatin accessibility that are correlated with cell type-specific changes in gene expression. Notably, regions with increased accessibility are enriched for AP-1 binding sites, suggesting a central role for the Fos family TFs in orchestrating these gene expression changes in neurons. Although Fos is a commonly induced TF across many cell types, we find that its chromatin binding sites and associated late response genes exhibit notable cell type specificity. This raises an open question about how a widely expressed TF like Fos can drive such distinct transcriptional programs in different cell types. Our findings therefore represent a rich resource to enable the functional characterization of the cell-type-specific LRGs that drive long-term structural changes in the hippocampus following NE exposure, as well as the *cis*-regulatory elements that drive these cell type-specific gene expression changes.

This resource contributes to a more comprehensive understanding of the early molecular response to naturalistic stimuli in the hippocampus. While substantial progress has been made in generating atlases of cell-type-specific gene expression and chromatin state.^27,49^ these previously collected datasets largely reflect conditions in naïve animals, thereby missing transient gene expression programs induced by salient environmental stimuli. Our study thus provides an important resource for future investigation into the functional consequences of these gene expression programs in the context of experience-dependent plasticity and learning.

For example, we identified specific ligand-receptor pairs whose expression is induced in response to novel environment exposure and which may act in a coordinated fashion across cell types. Ccn1, a secreted factor upregulated in CA1, CA3, and DG, is notable for its role in activating microglia—a cell type that is essential for clearing debris and promoting synaptic plasticity but which requires precise temporal control to avoid prolonged activation linked to neurodegeneration. Similarly, *Bdnf* and *Ntf3*, which bind to receptors expressed in multiple hippocampal cell types, may facilitate broad neuronal survival and synaptic remodeling, supporting learning and memory functions. These ligand-receptor interactions highlight a mechanism by which excitatory neurons may regulate neighboring cell responses, potentially shaping microenvironmental conditions in ways that support circuit-specific plasticity. Our identification of NE-induced ligands and their cell type-specific receptors therefore will enable researchers to dissect the precise roles these signaling molecules play in coordinating cell-cell interactions during spatial memory formation.

Future researchers can leverage this resource to strategically design functional studies on the relationship between NE exposure, chromatin opening, and gene expression in specific cell types. For example, our searchable database accompanying this manuscript allows researchers without prior computational experience to search for genes of interest and identify key cell types and time points of NE exposure when designing their experiments (**Fig. S1C**). Furthermore, recent advances in technologies that enable the high-throughput interrogation of *cis*-regulatory element activity in specific cell types^50–52^ could be used to clarify the contribution of these genomic elements to cell-type-specific NE-driven gene expression programs. Notably, the rapid activation of AP-1-related transcriptional programs in response to sensory stimuli appears to be a conserved mechanism across brain regions^5^ and species,^53,54^ suggesting that this ERG program is fundamental to a wide range of behavioral and cognitive processes.

Certain limitations of the study should be considered when interpreting the results. First, the microdissection of hippocampal regions for bulk RNA-seq provides a heterogeneous mix of cell types, and does not differentiate between mRNA localized to the nucleus, cytoplasm, dendrites or axons. While snMultiome-seq offers complementary information about gene expression and chromatin accessibility within specific cell types, this method captures nuclear RNA, making it difficult to detect genes that are transiently transcribed and rapidly exported to the cytoplasm, including many ERGs. This limitation could result in the underrepresentation of these genes in our dataset despite their robust transcriptional activity.

In summary, we present a comprehensive dataset characterizing the dynamic chromatin accessibility and gene expression landscape following acute NE exposure. The accompanying user-friendly database also provides a valuable resource for further exploration of activity-dependent genes and putative gene regulatory elements that will be critical to advance tool development for the targeted manipulation of AP1-dependent gene expression programs that contribute to novel environment encoding, as well as learning and memory.

## Author Contributions

L.T., E.E.D., and M.E.G. conceptualized the study and designed the experiments. E.E.D., L.T., and N.N. performed behavioral assays. L.T. performed hippocampal dissections. E.E.D., L.T., E.G.A., N.P. and N.N. performed bulk RNA-seq assays. S.Sun and H.L. performed snMultiome-seq assays. E.E.D., L.T. and H.L. performed bioinformatic analyses. S.Sanalidou performed immunohistochemistry and microscopy. E.E.D., L.T., E.C.G. and M.E.G. drafted the manuscript, with input from all co-authors.

## Acknowledgments

The Greenberg Laboratory is supported by the Allen Discovery Center Program, a Paul G. Allen Frontiers Group advised program of the Paul G. Allen Family Foundation and the Tang-Yang Autism Center at Harvard Medical School. L.T. was supported by the Long-term Human Frontiers Science Program Fellowship and the William Randolph Hearst Fund. E.E.D. was a Damon Runyon-National Mah Jongg League, Inc. Breakthrough Scientist supported by the Damon Runyon Cancer Research Foundation and was supported by the Warren Alpert Distinguished Scholar Award and a Mahoney Postdoctoral Fellowship. H.L. was supported by the Harvard Society of Fellows. M.E.G. was supported by funding from NINDS R01 NS115965. The funders had no role in study design, data collection and analysis, or the decision to publish or preparation of the manuscript. We thank members of the Greenberg laboratory for helpful discussions on the manuscript. We thank the Harvard Medical School Neurobiology Department for consultation and instrument availability that supported this work.

## Data availability

All bulk RNA and snMultiome sequencing data have been deposited in the Gene Expression Omnibus database under accession number GSE283483 (reviewer access key: irwxkkkcfnubnsx). Our web-based searchable database is available from (https://greenberg.hms.harvard.edu/project/novel-environment-gene-database/).

## Code availability

All code used in this study was previously published and is cited in the Methods section.

## METHODS

### Mice

Mice were handled according to protocols approved by the Harvard University Standing Committee on Animal Care and were in accordance with federal guidelines. Wild-type C57/BL6 (JAX 000664) were housed in a 12 h light/dark cycle with ad libitum access to food. For all sequencing studies, male mice were used to avoid the confounding effects of estrogen cycles observed in females, but for histology experiments female mice were used. While staining for ERGs in female mice yielded similar results (**Fig. 1B, S1A**), supporting the relevance of these pathways across sexes, this approach limits our ability to directly translate our findings to both sexes.

### Novel environment paradigm

Adult mice (8-10 weeks of age) were single housed and brought to the room in which behavior was conducted 2-3 nights prior to exposure to novel environment. During this time, mice were housed in a 12 h light/dark cycle with ad libitum access to food. For novel environment stimulation, mice were placed in a square arena (16”x16”) containing a running wheel, tunnel, falcon tubes, Lego pyramid, and a tube rack for 30 min, returned to their home cage, and brain tissue was harvested at indicated times after the start of novel environment (except for 15 and 30 min timepoints for which tissue was harvested without return to homecage). For 6 h continuous novel environment stimulation, mice were placed into the arena containing toys and hydro gel for 6 h. Arena and toys were cleaned prior to experimentation and in between animals with 70% EtOH. Importantly, tissue was always collected before the animals entered the dark cycle, so the gene expression changes observed are likely not activated in response to changes in response to light state. We collected replicates from the same time point at different times of day to control for gene expression changes driven by circadian rhythms rather than NE exposure. Our initial dataset also included bulk RNA-seq following 12 h of NE exposure, but this time point was later excluded as all mice were harvested at the same time immediately before the dark cycle began and therefore gene expression changes due to NE exposure and circadian rhythms could not be deconvoluted.

### Microdissection of hippocampus

Mice were deeply anesthetized with isoflurane, decapitated and brains were placed in a stainless-steel brain matrix on ice to cut 1mm sections of anterior to posterior hippocampus. Sections were transferred into ice-cold PBS and Cornu Ammonis (CA)1, CA3 and Dentate Gyrus (DG) separated under a dissection microscope. Tissue was frozen in liquid nitrogen and stored at -80° for further processing and RNA extraction. For single-nuclei multiomic-sequencing whole hippocampi were dissected on an ice-cold metal block and immediately processed for nuclei preparation (see below).

### RNA isolation and library preparation for bulk RNA-seq

Tissue was flash-frozen and resuspended in 500 µL TRIzol reagent (Life Technologies), followed by trituration with a 26G needle. RNA was chloroform extracted, an equal volume of 100% ethanol was added, and RNA was purified using the RNeasy Micro Kit (Qiagen) according to the manufacturer’s instruction, including the on-column DNase digest. RNA concentration was assessed by Nanodrop and RNA-seq libraries were prepared from 10 ng of total RNA using the SMARTer Stranded Total RNA-seq Pico Input Mammalian V2 kit (Takara Bio) according to the manufacturer’s instructions. Samples were multiplexed with Illumina TruSeq HT barcodes and sequenced on a NovaSeq X 25B with paired-end 2 × 150-nt reads. All samples were sequenced to a depth of at least 20M reads.

### Bulk RNA-seq Analysis

Illumina adapters were trimmed with Cutadapt (version 1.14) and aligned using Hisat2^56^ (version 2.1.0) to the *Mus musculus* genome (GRCm39) and transcriptome (Ensembl). Alignments and analysis were performed on the Orchestra2 high performance computing cluster through Harvard Medical School. Aligned BAM files were sorted using Picard Tools (version 2.8.0); stranded bedGraphs were generated using STAR^57^ (version 2.7.0f); and reads were quantified over annotated exons using FeatureCounts^58^ (Subread version 2.0.6). Lowly expressed genes were filtered for counts per million >1 in at least five samples using edgeR (version 4.0.16).

Differential expression analysis was performed using DESeq2.^59^ Time points from the same region were analyzed together and both time of NE exposure and library preparation batch effects were considered in the design (∼ time + batch). Fold change shrinkage was applied using apeglm.^60^ Significant genes were defined as p_adj_ < 0.05 and abs(log_2_(fold change)) > log_2_(1.2) after shrinkage. For HC gene expression comparisons in bulk RNA-seq, DESeq2 was run on pairwise comparisons (CA1 vs. CA3, CA1 vs. DG, CA3 vs. DG) and fold change shrinkage was applied. Significant genes were defined as p_adj_ < 0.05 and abs(log_2_(fold change)) > log_2_(2) after shrinkage. Depth-normalized counts were generated using DESeq2. Gene Ontology (GO) enrichment analysis was performed using gProfiler2 in R (version 0.2.3), with a custom background of expressed genes based on expression-filtered RNA-seq genes and false discovery rate (FDR) < 0.05. Heatmaps were generated using ComplexHeatmap (version 2.18.0).^61^ TFs and secreted factors were defined by the GO terms for DNA-binding transcription factor activity (GO:0003700) and secretion (GO:0046903), respectively. Synaptic genes were defined using the SynGO database.^62^ Pairs of secreted factors with known receptors were identified using CellTalkDB (http://tcm.zju.edu.cn/celltalkdb/).

### Single-cell deconvolution from bulk RNA-seq data

The SCDC R package (version 0.0.0.9000)^55^ was used to approximate the distribution of cell types in our bulk RNA-seq, using single-cell RNA-seq data from the Allen Brain Atlas as a cell type reference.^27^ Raw counts data from samples in the reference dataset were normalized, and the distribution of cell types present in each of the bulk RNA-seq samples in this study was determined using SCDC.

### Single-nuclei Multiome nuclei preparation

Freshly dissected hippocampi were dissociated, and nuclei were prepared as previously described.^63^ Nuclei were then concentrated in 1X nucleus buffer from 10X Genomics. 1µl of nuclei was stained with Trypan Blue (Invitrogen, T10282) and manually counted. In total, 16,000– 20,000 nuclei were used for the tagmentation reaction and 10x Chromium controller loading. The DNA and RNA libraries were generated according to the manufacturer’s recommended protocol (https://www.10xgenomics.com/support/single-cell-multiome-atac-plus-gene-expression). 10x multiome ATAC–seq and RNA-sequencing (RNA-seq) libraries were sequenced together on the Illumina NovaSeq X systems using paired-end 2x150bp reads to a depth of around 50,000 reads per cell for each modality.

### snRNA-seq Analysis

Raw sequencing data were processed using Cell Ranger ARC (v2.0.2, 10x Genomics) under default settings. Reads were mapped to mouse mm10 genome, with the GENCODE vm23 GTF file as used in the BICCN atlas.^27^ We performed unsupervised clustering with RNA UMI counts based on the SCANPY analysis pipeline^64^ and best practice steps.^65,66^ In brief, we first filtered for low-quality nuclei by requiring ≥1,000 ATAC fragments and ≥200 genes detected per nuclei and < 5% mitochondria RNA reads. We then performed a first-pass clustering for quality control. Top 3,000 highly variable genes were identified for principal component analysis (PCA). Top 30 PCs were used to perform K-nearest neighbor (k=25) graph-based Leiden clustering (resolution=1). Putative multiplets were predicted using Scrublet^67^ and Leiden clusters with mean doublets score > 0.2 were marked as doublets. 87-92% cells generated from Cell Ranger ARC for each replicate passed these filtering steps. We then integrated our high-quality cells with hippocampal formation (HPF) cells selected from the whole-mouse brain atlas dataset from BICCN.^27^ The integration was performed using Seurat’s integration methods as previously described.^68^ We annotated 14 major cell types, and 33 sub-class level cell types based on the integration results as the cell type nomenclature used in this study. To visualize clusters and annotations, we used scVI package^69^ (v1.2.0 n_hidden = 512, n_latent = 40, n_layers = 2, use sample as batch_key) to learn a latent space and perform UMAP embedding^70^ to generate the visualization coordinates for scatter plots.

### snATAC-seq analysis

We used SnapATAC2 (2.5.0)^71^ to process fragment files generated from Cell Ranger ARC to make the cell-by-peak h5ad file and bigwig files for each cell type. We used the PoissonVI model^72^ from the scVI-tools^69^ to learn a cell-by-peak based latent embedding and performed differential peak accessibility analysis. This analysis is done on peaks accessible in >1% of all cells. Differential accessible peaks based on the PoissonVI model were filtered by FDR < 0.05, lfc_mean > 0.3 and emp_lfc > 0.3. For Chromvar analysis,^48^ we used the whole cell-by-peak matrix as input and scanned motifs across the JASPAR2024 motif database.^73^ We then performed a two-sided t-test with Benjamini-Hochberg FDR correction on the cell-level Chromvar scores and selected significant differential motifs based on an FDR < 0.05. Similar motifs belonging to the same motif family were merged and the most differential motif was selected to represent the motif family (e.g Fos for the AP-1 family). To test whether the variance of cell-level gene expression can be explained by a motif’s Chromvar score (e.g., Fos), we grouped cells into five equal-sized bins based on their Chromvar score and performed a one-way ANOVA test with FDR correction using the Pingouin package (0.5.5).^74^ For genes with significant ANOVA results (FDR < 0.05), we further performed a post-hoc Tukey pairwise test to determine the relationship between the Chromvar score and gene expression value. For this analysis, we used log_10_ (1 + transformed count per million (CPM)) values for the gene expression. For Homer (v4.11)^75^ motif analysis, we used default Homer motif datasets and selected the top 3000 most differentially accessible peaks based on their accessibility fold change between two conditions and resized them into 500bp bins. We then performed findMotifsGenome.pl and used one condition as foreground (1h NE and 6h cNE) and the other as background (HC) to test for motif enrichment. Differential expression analysis was performed on the cell level for a given cell type, comparing the distribution of expression between HC and NE, using the Wilcoxon rank-sum test for statistical significance, followed by multiple hypothesis correction with Bonferroni adjustment.

### Histology

Mice were deeply anesthetized with 9 mg/kg ketamine-xylazine by intraperitoneal injection, and transcardially perfused with ice-cold PBS followed by 4% paraformaldehyde (PFA) in PBS. Dissected brains were post-fixed overnight at 4°C, next incubated overnight in 15% sucrose prepared in PBS, and overnight in 30% sucrose in PBS at 4°C. Brains were sectioned on a vibratome LeicaVT1000 at 40µm. Free floating hippocampal sections were permeabilized with 0.25% Triton in PBS 1X for 1 h, and blocked (5% bovine serum albumin, 0.3% Triton 100X, and 10% donkey serum in PBS 1X) for 2 h. Blocking solution was replaced by primary antibody solution (0.1% Triton 100X and 10% donkey serum in PBS 1X). After an overnight incubation at room temperature with the primary antibodies: Egr1 anti-rabbit (1:2000, Cell Signaling Technology #4153S) and c-Fos anti-guinea pig (1:3000, Synaptic Systems, #226308), brain slices were washed 3x in 0.1% Triton 100X in PBS for 15 min each. Sections were incubated in secondary antibody solution (0.1% Triton 100X in PBS) for 2 h at room temperature with the following secondary antibodies: Cy3 donkey anti-rabbit IgG (1:1000, Jackson ImmunoResearch, #711-165-152) and Cy5 donkey anti-guinea pig IgG (1:1000, Jackson ImmunoResearch, #706- 175-148). Slices were washed twice at RT with 0.1% Triton 100X in PBS and then counterstaining with 4’,6-diamidine-2’-phenylindole dihydrochloride (DAPI) (1:5000, Sigma Millipore, #MBD0015) in PBS for 5 min. Finally, brain slices were washed twice in 1X PBS. Images were acquired with 10X and 40X objectives, at 1024 x 1024 pixel resolution, and 16-bit depth on a ZEISS LSM800 confocal microscope.

## TABLES

Table 1: Fold changes and adjusted p-values for all comparisons in bulk and snRNA-seq

Table 2: GO analysis enriched terms

Table 3: Chromvar scores for each gene and cell type

---

**Extended Data Figure 1:**
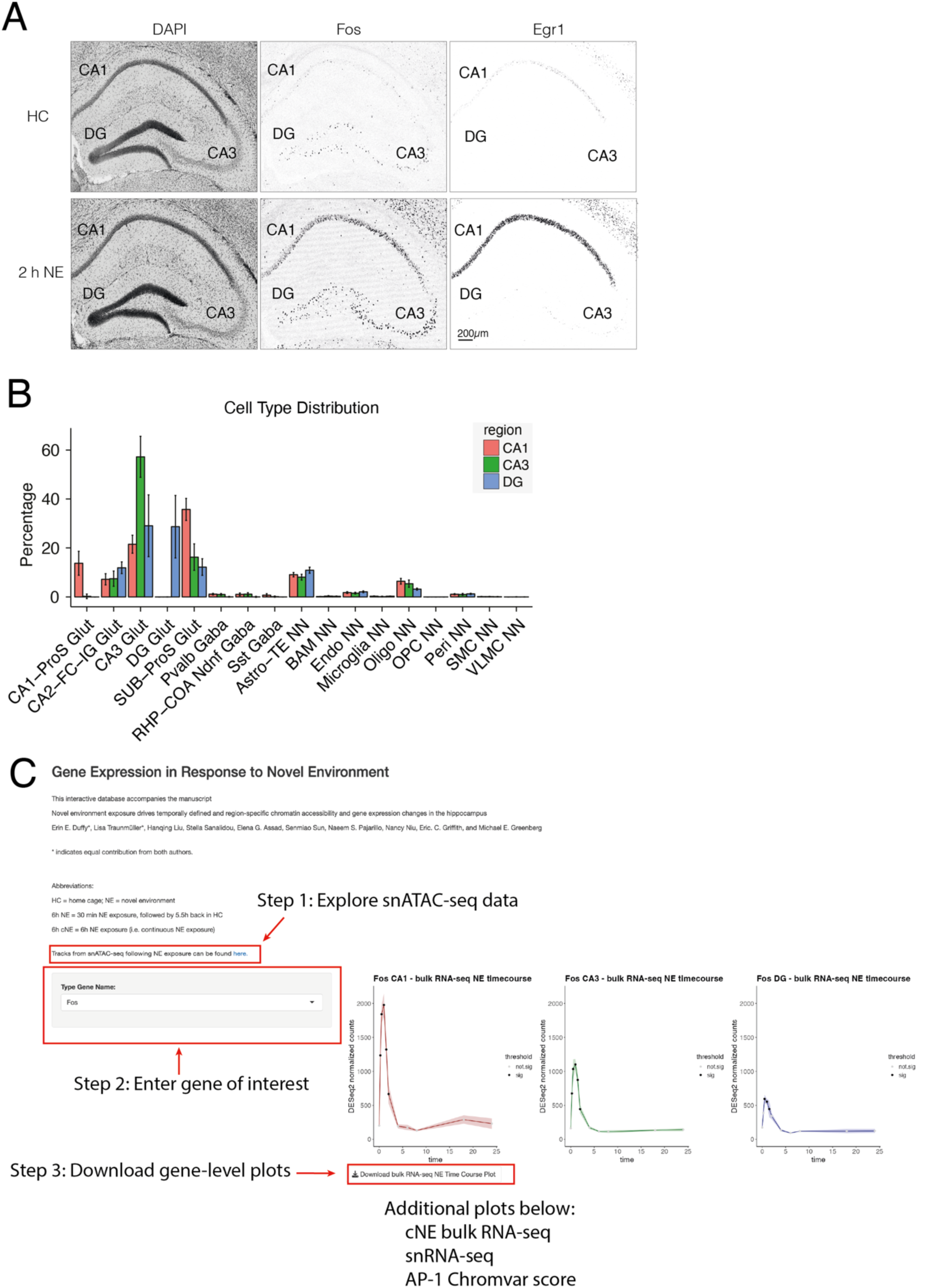
NE exposure stimulates hippocampal ERG activity, Related to. Figure 1**. (A)** Immunofluorescence images of FOS, and EGR1 protein levels in HC and 2 h NE (30 min NE + 1.5 h HC) in CA1, CA3, and DG. Scale bar 200 µm **(B)** As differences in cell type composition can drive differential expression in bulk RNA measurements, we used single-cell deconvolution^55^ to approximate the proportion of neuronal and non-neuronal cells in each sample by region, using the Brain Initiative Cell Census Network 2.0^27^ as a reference. Bar plot of bulk gene expression deconvolution of RNA-seq from CA1, CA3, and DG across all time points of NE exposure. Differences in non-neuronal cell type composition are not statistically significant by two-way ANOVA (F statistic. = 0.00, P=1.000), strongly suggesting that observed differences between regions are driven primarily by gene expression changes in excitatory cells rather than non-neuronal cells. Data are shown as mean ± SD, n=48-50 biologically independent tissue samples per region. **(C)** Tutorial on using the online database. Users can explore snATAC-seq tracks in the UCSC genome browser, enter a gene of interest, and download high-resolution plots of bulk RNA-seq expression following brief and cNE exposure, as well as cell type-specific gene expression (snRNA-seq) and AP-1 Chromvar scores.

**Extended Data Figure 2:**
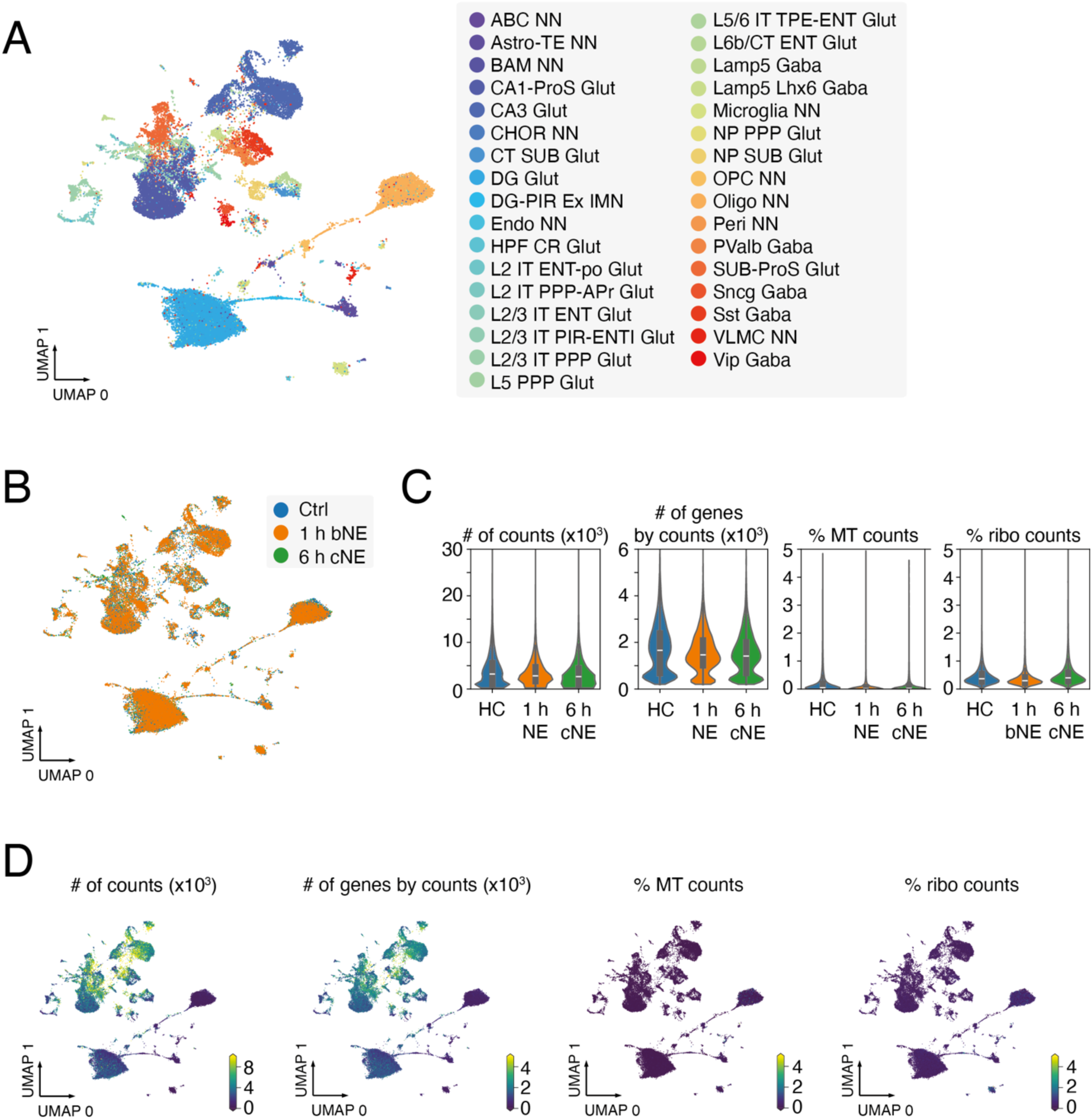
snRNA-seq quality metrics, Related to. Figure 1**. (A)** UMAP visualization of nuclei from HC, 1 h brief NE (bNE), and 6 h continuous NE (cNE) snRNA-seq with cell sub-type information overlaid. n = 45,844 cells, 3 mice. **(B)** UMAP as in (A), with sample information overlaid. **(C)** Violin plot of number of counts per cell, number of genes by counts, percentage of mitochondrial RNA detected per cell (% MT counts), and percentage of ribosomal RNA detected per cell (%ribo counts), separated by sample (HC, 1 h brief NE, 6 h cNE). **(D)** UMAP as in (A), with information from C overlaid.

**Extended Data Figure 3:**
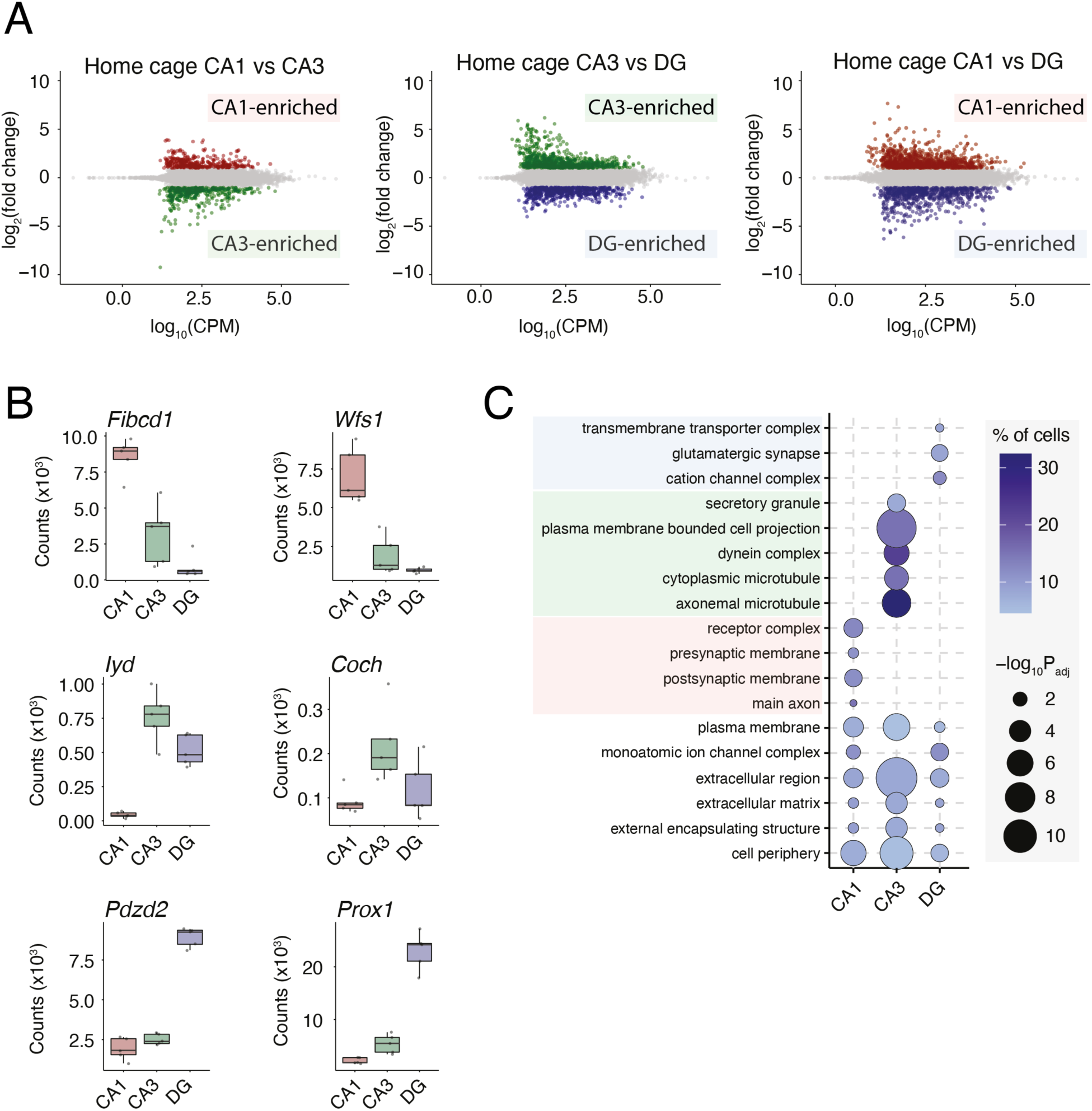
Region-specific differences in HC gene expression, Related to. Figure 2**. (A)** MA plot of pairwise differentially expressed genes in HC condition between CA1 (red), CA3, (green) and DG (blue). Colored points represent significant (DESeq2 P_adj_ < 0.05) enrichment in one region. (**B**) Box and whisker plots of HC bulk RNA-seq depth-normalized counts for genes defined as region-specific marker genes by Cembrowski et al. 2016.^26^ **(C)** Dot plot of the top enriched GO terms for CA1- CA3- and DG-enriched genes in Figure 2A, as well as GO terms that were common between all three cell classes.

**Extended Data Figure 4:**
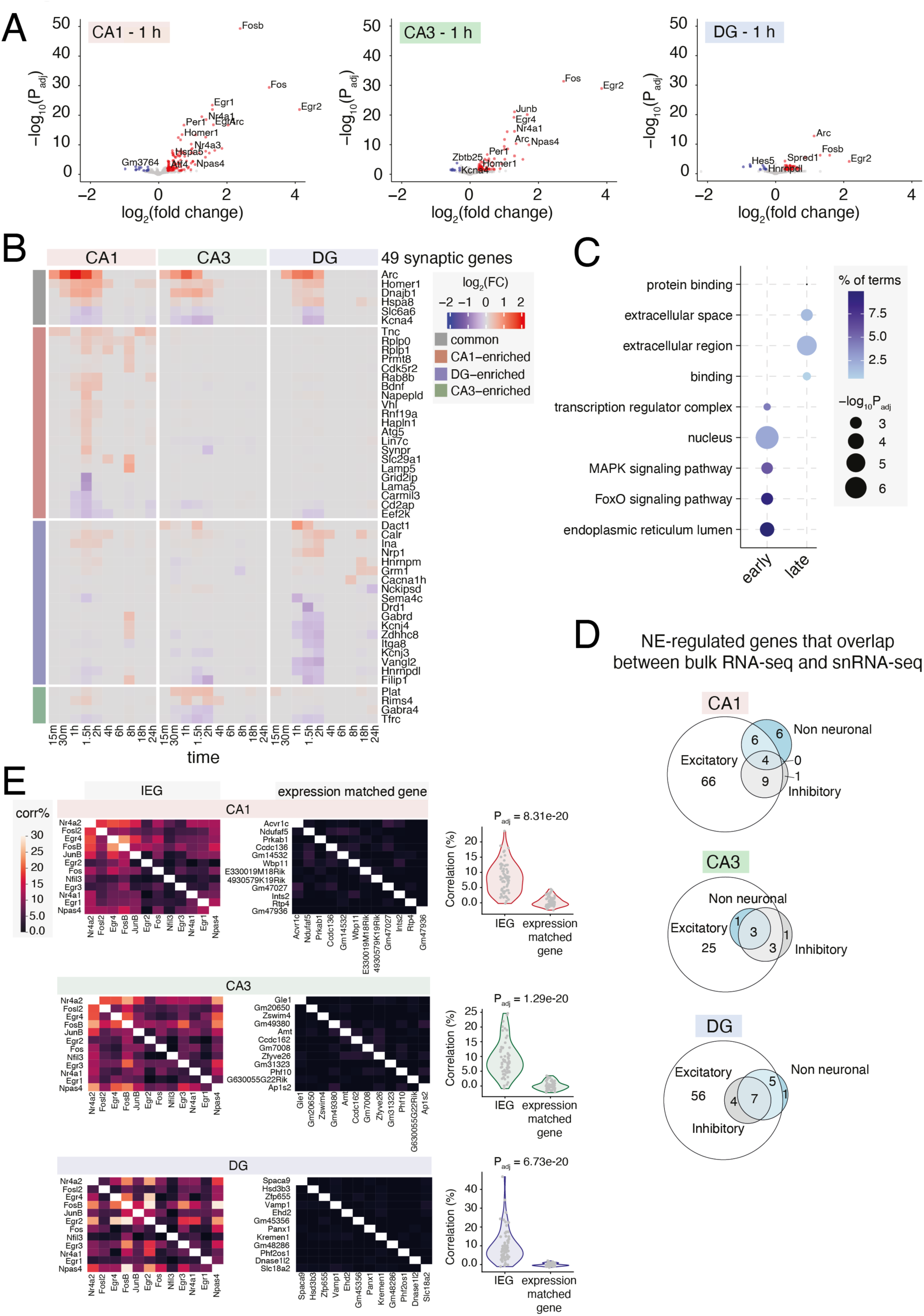
NE-driven gene expression changes, Related to. Figure 3**. (A)** Volcano plot of −log_10_(*P*_adj_) versus log_2_(fold change) in bulk RNA-seq expression between 1 h brief NE exposure and HC in CA1, CA3, and DG. Red indicates upregulated genes (DEseq2 *P*_adj_ < 0.05, Log_2_ fold change > log_2_(1.2)); blue indicates downregulated genes (DEseq2 *P*_adj_ < 0.05, Log_2_ fold change < -log_2_(1.2)); and gray indicates non-significant genes (DEseq2 *P*_adj_ > 0.05). **(B)** Heatmap as in (Fig 3C) for genes that are annotated as synaptic genes. **(C)** Dot plot of the top GO terms in early and late upregulated genes by bulk RNA-seq. ERGs were enriched for GO terms related to transcription factor activity in the nucleus, whereas late response genes were enriched for terms related to extracellular secretion and receptor binding. **(D)** Venn diagrams of genes detected as differentially expressed in both bulk and snRNA-seq. Genes were classified as CA1 CA3 or DG based on differential expression in bulk RNA-seq and further classified as excitatory, inhibitory, or non-neuronal based on the cell type(s) in which the same gene was differentially expressed in snRNA-seq data at 1 h NE. **(E)** Pairwise Pearson correlations across individual excitatory neurons in CA1, CA3, or DG at 1 h NE exposure, calculated on the basis of (left) commonly induced TF expression (Fig. 3D), or (right) expression-matched non- induced genes. Correlations between ERG TFs are significantly higher than those between expression-matched noninduced genes (p-values as indicated, Mann-Whitney *U* test, two-sided).

**Extended Data Figure 5:**
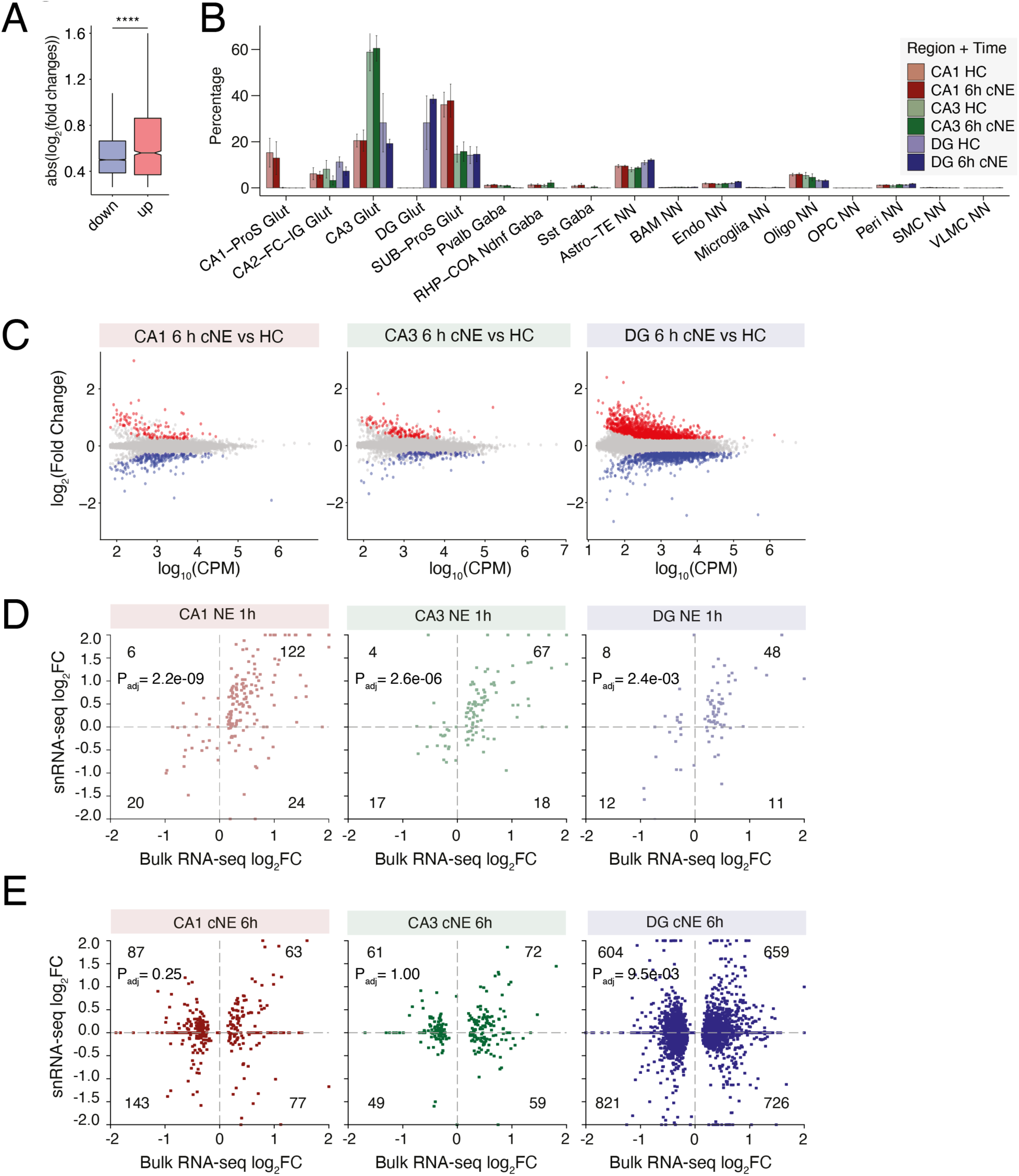
Continuous NE exposure boosts expression of LRGs, Related to. Figure 6**. (A)** Box plot of absolute value(log_2_ fold changes) in NE-upregulated (red) and - downregulated (blue) genes by snRNA-seq in Figure 5A. **** = P < 2.2e-16, Welch’s Two Sample t-test. **(B)** Bar plot of bulk gene expression deconvolution of RNA-seq from CA1, CA3, and DG, separated by HC and cNE exposure, using the Mouse Brain Atlas as a reference. Differences in cell type composition are not statistically significant between HC and cNE for each region by two-way ANOVA (F statistic = 0.048, P-value = 1.000), strongly suggesting that observed gene expression changes are driven by exposure to cNE and not cell type differences. Data are shown as mean ± SD, n=4-5 biologically independent replicates per region and time. **(C)** MA plot of pairwise differentially expressed genes between HC and 6 h cNE in CA1, CA3, and DG. Red indicates upregulated genes (DEseq2 *P*_adj_ < 0.05, FC > 1.5); blue indicates downregulated genes (DEseq2 *P*_adj_ < 0.05, FC < 0.667); and gray indicates non-significant genes (DEseq2 *P*_adj_ > 0.05). **(D & E)** Scatterplot of log_2_FC between bulk RNA-seq (x-axis) and snRNA-seq (y-axis) for genes defined as significant (DESeq2 Padj < 0.05) in bulk RNA-seq within region (CA1, CA3, DG) and time point (D = 1 h brief NE, E = 6 h cNE). P_adj_ = Pearson’s R correlation.

**Extended Data Figure 6:**
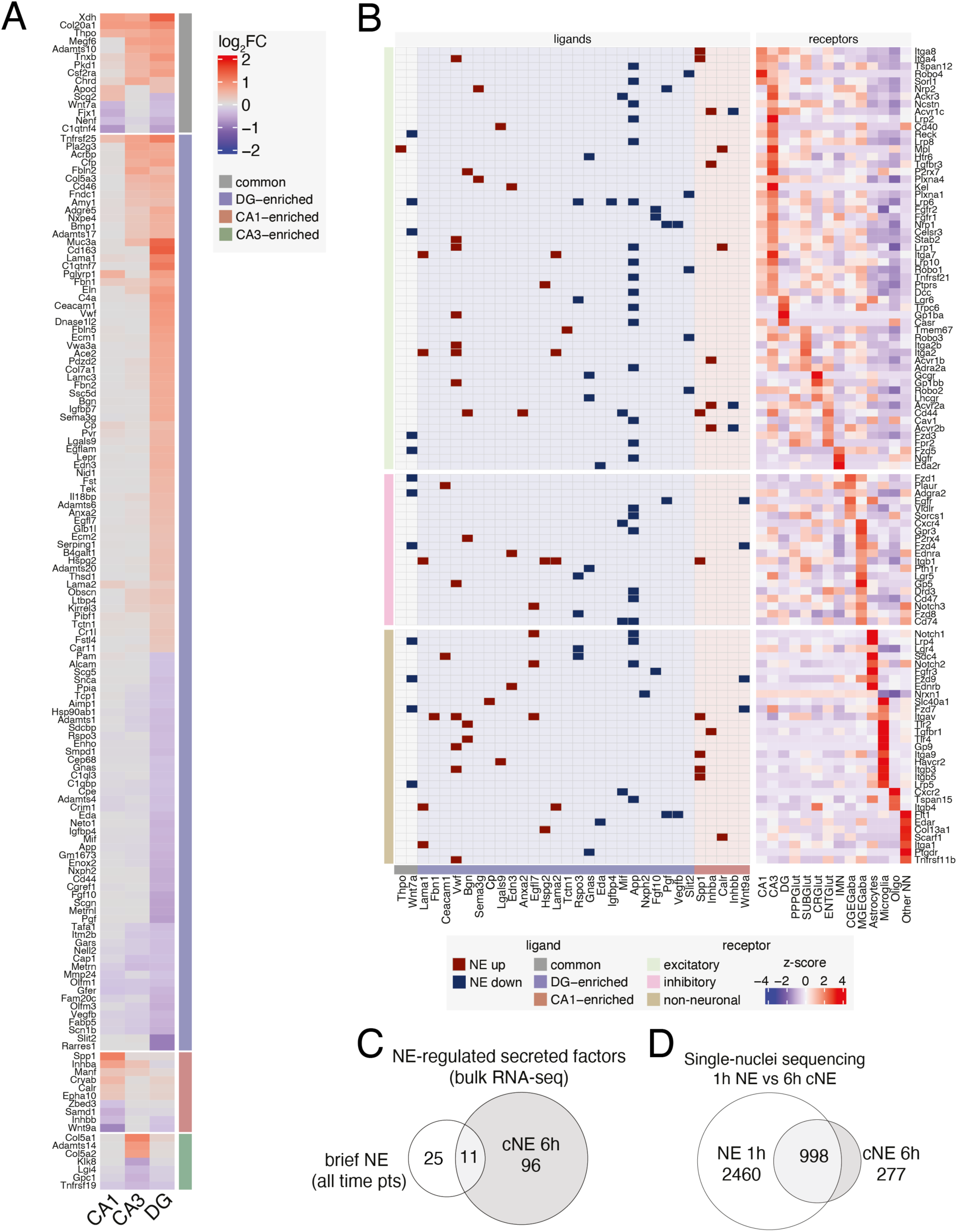
Secreted factors induced by 6 h cNE and their receptors, Related to. Figure 6**. (A)** Heatmap of log_2_(fold change) in gene expression of all significant NE-regulated genes that are annotated secreted factors at 6 h cNE versus HC by bulk RNA-seq. Colors over gene names indicate that the gene has at least one known receptor. **(B)** Interaction grid of NE-regulated secreted factors (ligands) and their receptors. Left grid shows ligand-receptor pair, with dark shading indicating whether ligand expression was upregulated (dark red) or downregulated (dark blue) in response to NE. Background shading indicates the hippocampal region of ligand NE gene expression regulation (common, CA1- CA3- or DG-enriched). Right heatmap shows row- normalized z-scores of cognate receptor expression from HC snRNA-seq across all major cell types, ordered by cell type of maximum receptor expression. (**C**) Venn diagram of secreted factors that are NE-induced following brief (all time points) or 6 h cNE exposure as displayed in Fig4C and S6A, respectively. (**D**) Venn diagram of all differentially regulated genes in snRNA-seq between 1 h brief and 6 h cNE.

**Extended Data Figure 7:**
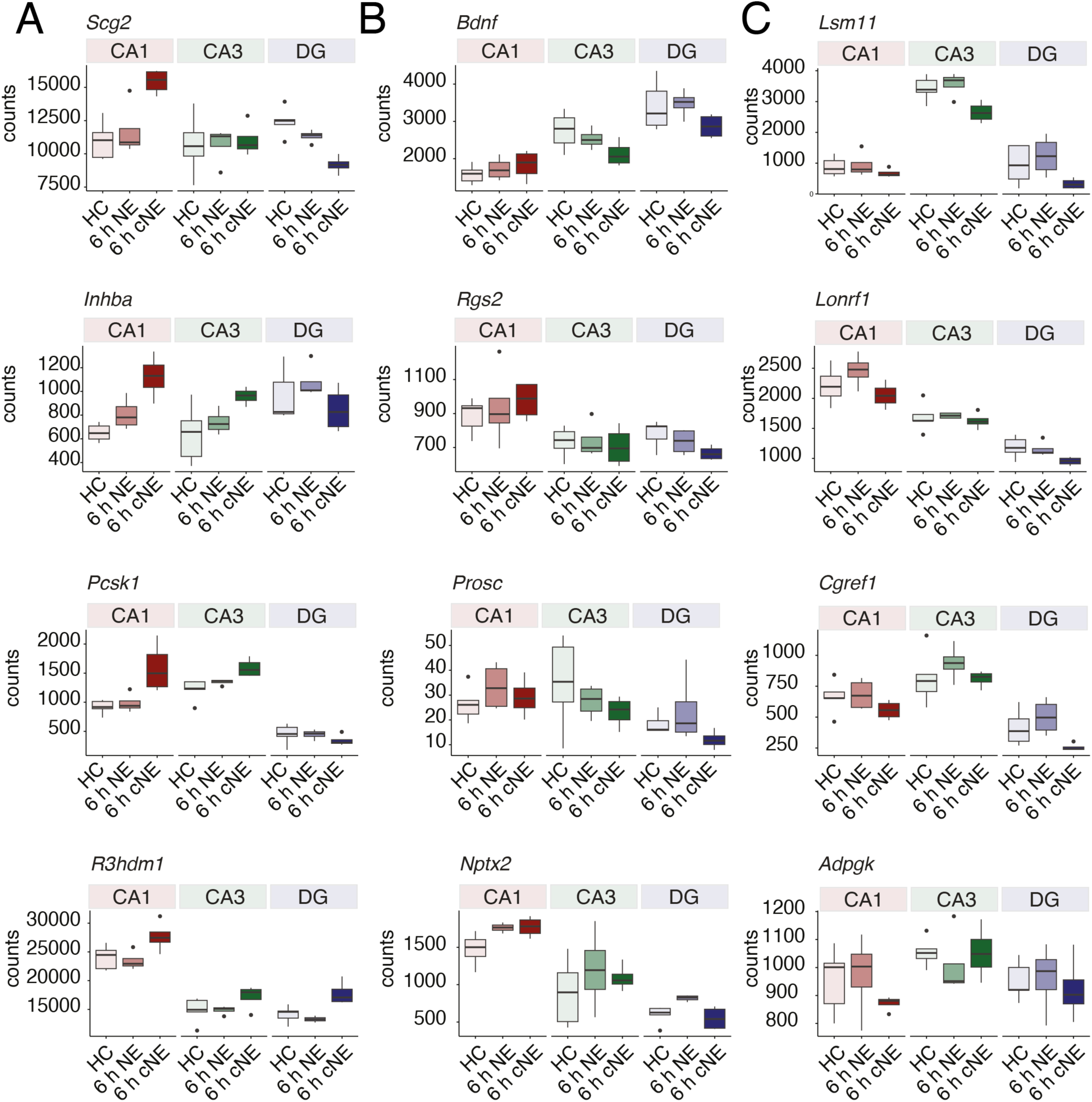
Gene expression changes of AP-1 targets following NE exposure, Related to. Figure 6. Box and whisker plots of bulk RNA-seq depth-normalized counts of gene expression in HC and following brief and cNE exposure for genes previously reported to be activated by Fos/AP-1 in response to kainic acid-induced seizures in CA1.^6^ Box plots shown as median ± IQR (whiskers = 1.5*IQR).

**Extended Data Figure 8:**
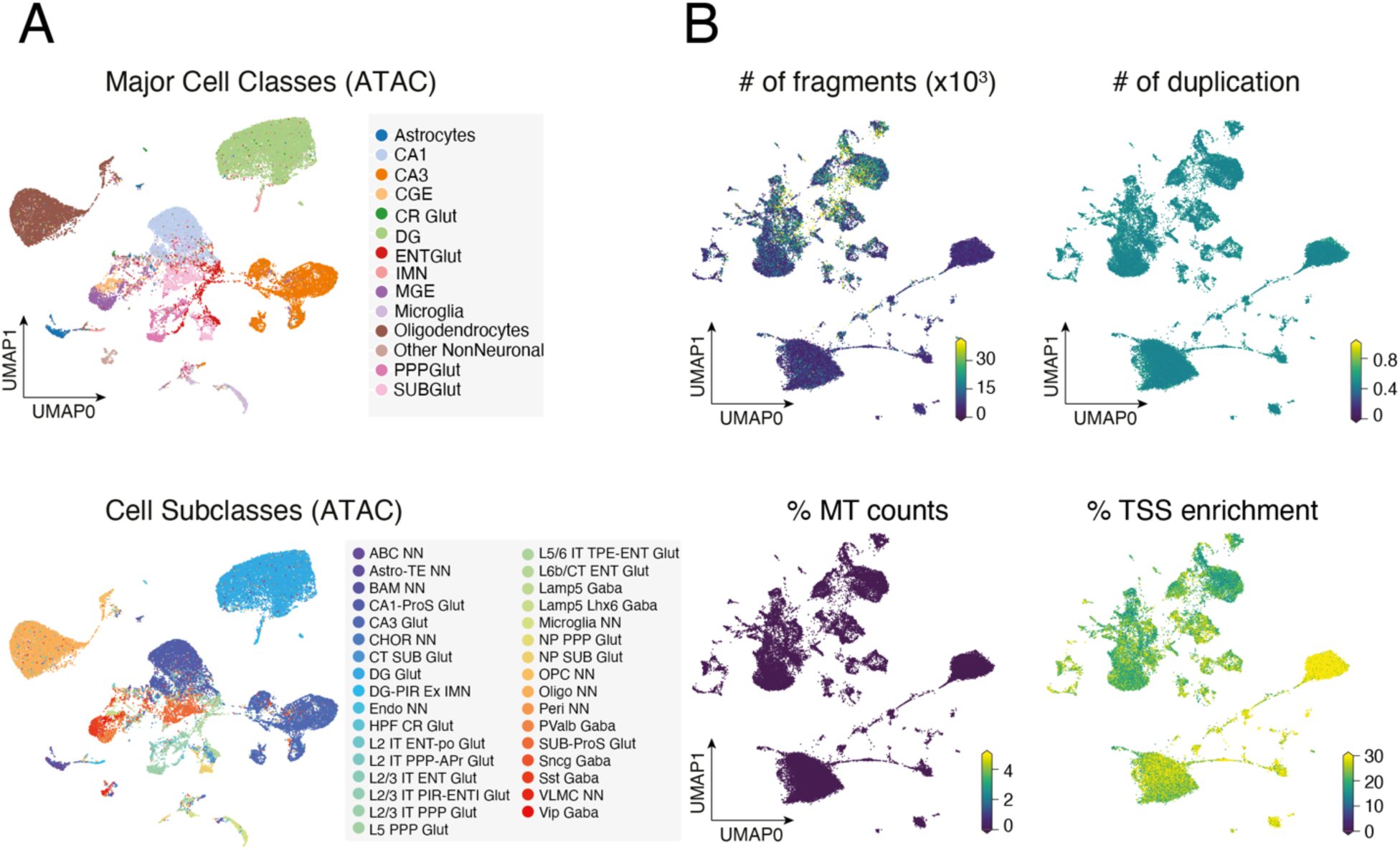
Chromatin accessibility changes in response to NE at single-cell resolution, Related to. Figure 7**. (A)** UMAP visualization of nuclei from HC, 1 h brief NE, and 6 h cNE snATAC-seq with cell type (top) and cell sub-type (bottom) information overlaid. n = 45,844 cells, 3 mice. **(B)** UMAP as in (Fig 1C), with number of fragments per cell, fraction of duplicated fragments per cell (# of duplication), fraction of mitochondrial fragments per cell (% MT counts), and % transcription start site enrichment (% TSS enrichment) information overlaid.

**Extended Data Figure 9:**
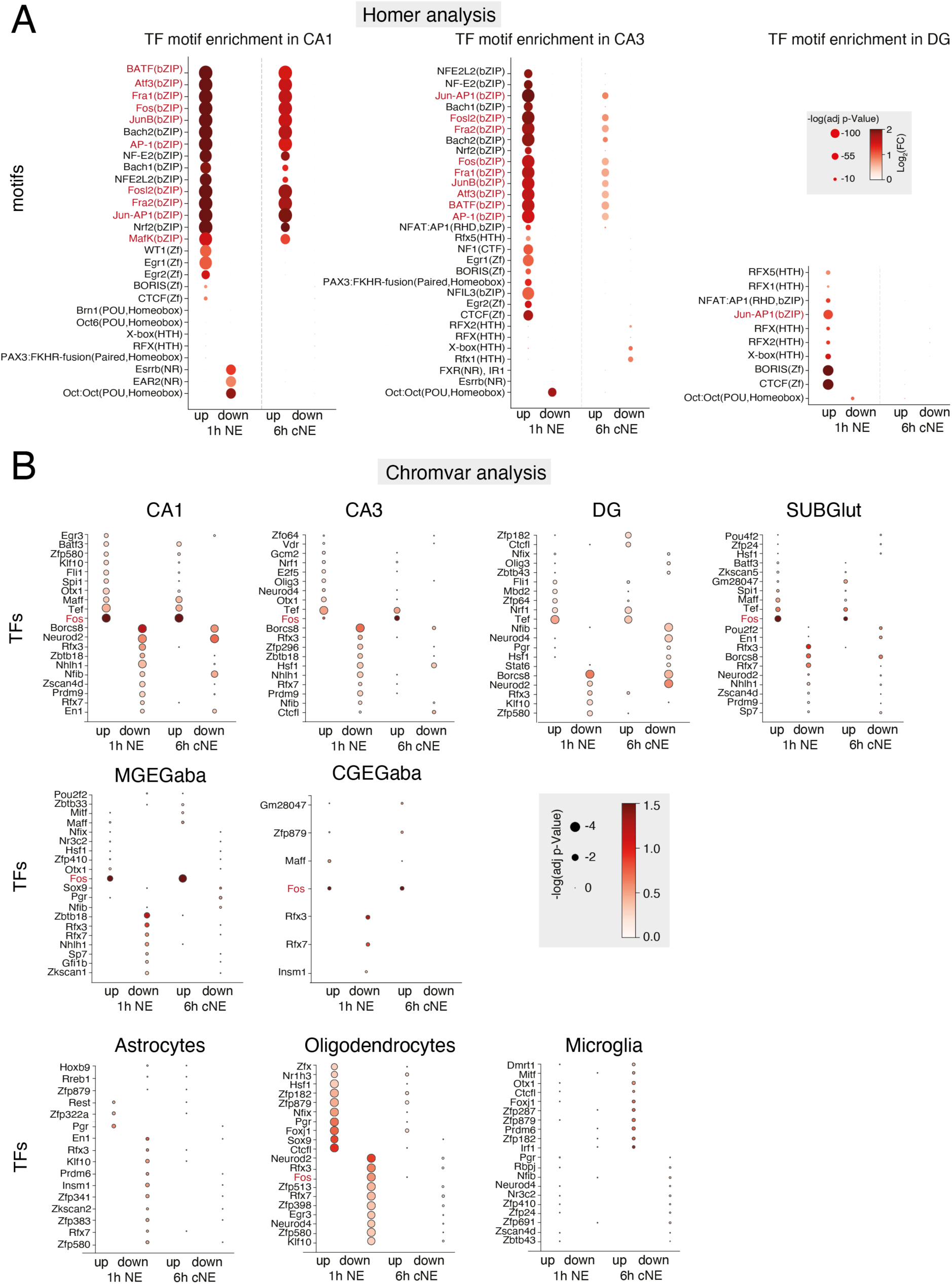
Motif enrichment in activity-regulated ATAC peaks, Related to. Figure 7**. (A)** Dot plot of TF motifs enriched in CA1, CA3, and DG excitatory neurons within differentially accessible peaks following 1 h brief NE or 6 h cNE compared to HC using Homer. AP-1-related motifs are indicated in red. **(B)** Dot plot of TF motifs enriched in major excitatory- and inhibitory neurons, and non-neuronal cells within differentially accessible peaks following 1 h brief NE or 6 h cNE compared to HC using Chromvar. Fos is indicated in red.

**Extended Data Figure 10:**
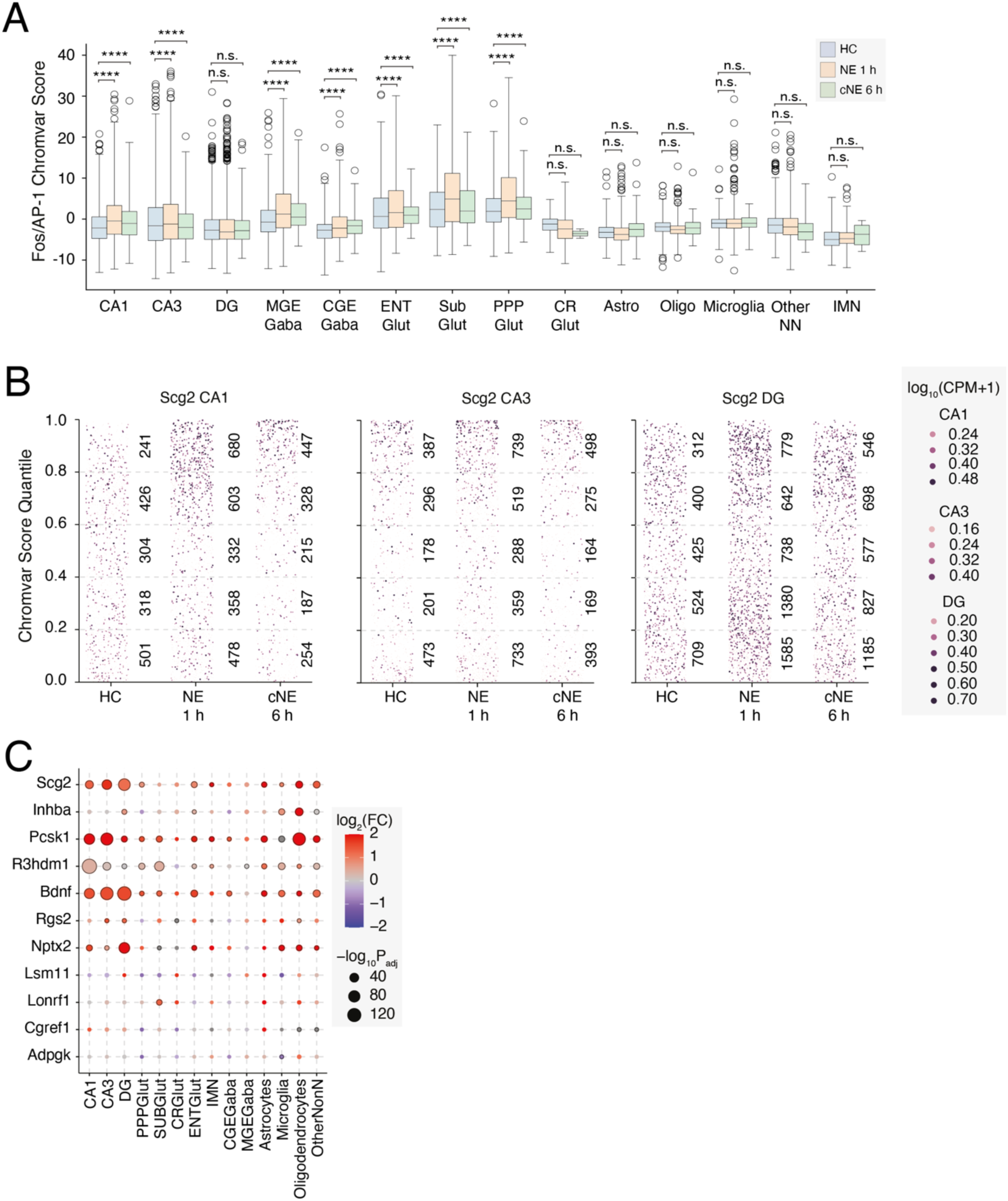
Motif enrichment in activity-regulated ATAC peaks, Related to. Figure 7**. (A)** Box and whisker plot of Fos/AP-1 Chromvar scores within all major cell types. (**B**) Dot plot of *Scg2* expression in CA1, CA3, and DG excitatory neurons. Cells are plotted based on Fos/AP-1 Chromvar score, where dotted lines indicate Chromvar score quantiles, numbers indicate the number of cells in each quantile, and color indicates *Scg2* expression in log_10_CPM. **(C)** Dot plot of log_2_FC in Fos/AP-1 Chromvar scores of genes displaying up-regulation in bulk RNA-seq after 6 h cNE, no change or down-regulation based on Fig.S7.

## References

1. Yap, E.-L., and Greenberg, M.E. (2018). Activity-Regulated Transcription: Bridging the Gap between Neural Activity and Behavior. Neuron 100, 330–348. 10.1016/j.neuron.2018.10.013.

2. Kaminska, B., Kaczmarek, L., and Chaudhuri, A. (1996). Visual Stimulation Regulates the Expression of Transcription Factors and Modulates the Composition of AP-1 in Visual Cortexa. J. Neurosci. 16, 3968–3978. 10.1523/JNEUROSCI.16-12-03968.1996.

3. Majdan, M., and Shatz, C.J. (2006). Effects of visual experience on activity-dependent gene regulation in cortex. Nat. Neurosci. 9, 650–659. 10.1038/nn1674.

4. Mardinly, A.R., Spiegel, I., Patrizi, A., Centofante, E., Bazinet, J.E., Tzeng, C.P., Mandel-Brehm, C., Harmin, D.A., Adesnik, H., Fagiolini, M., et al. (2016). Sensory experience regulates cortical inhibition by inducing IGF1 in VIP neurons. Nature 531, 371–375. 10.1038/nature17187.

5. Hrvatin, S., Hochbaum, D.R., Nagy, M.A., Cicconet, M., Robertson, K., Cheadle, L., Zilionis, R., Ratner, A., Borges-Monroy, R., Klein, A.M., et al. (2018). Single-cell analysis of experience-dependent transcriptomic states in the mouse visual cortex. Nat. Neurosci. 21, 120–129. 10.1038/s41593-017-0029-5.

6. Yap, E.-L., Pettit, N.L., Davis, C.P., Nagy, M.A., Harmin, D.A., Golden, E., Dagliyan, O., Lin, C., Rudolph, S., Sharma, N., et al. (2021). Bidirectional perisomatic inhibitory plasticity of a Fos neuronal network. Nature 590, 115–121. 10.1038/s41586-020-3031-0.

7. Lin, Y., Bloodgood, B.L., Hauser, J.L., Lapan, A.D., Koon, A.C., Kim, T.-K., Hu, L.S., Malik, A.N., and Greenberg, M.E. (2008). Activity-dependent regulation of inhibitory synapse development by Npas4. Nature 455, 1198–1204. 10.1038/nature07319.

8. Tyssowski, K.M., DeStefino, N.R., Cho, J.-H., Dunn, C.J., Poston, R.G., Carty, C.E., Jones, R.D., Chang, S.M., Romeo, P., Wurzelmann, M.K., et al. (2018). Different Neuronal Activity Patterns Induce Different Gene Expression Programs. Neuron 98, 530–546.e11. 10.1016/j.neuron.2018.04.001.

9. Spiegel, I., Mardinly, A., Gabel, H., Bazinet, J., Couch, C., Tzeng, C., Harmin, D., and Greenberg, M. (2014). Npas4 regulates excitatory-inhibitory balance within neural circuits through cell type-specific gene programs. Cell 157, 1216–1229. 10.1016/j.cell.2014.03.058.

10. Ataman, B., Boulting, G.L., Harmin, D.A., Yang, M.G., Baker-Salisbury, M., Yap, E.-L., Malik, A.N., Mei, K., Rubin, A.A., Spiegel, I., et al. (2016). Evolution of Osteocrin as an activity-regulated factor in the primate brain. Nature 539, 242–247. 10.1038/nature20111.

11. Ma, Y., Bendl, J., Hartley, B.J., Fullard, J.F., Abdelaal, R., Ho, S.-M., Kosoy, R., Gochman, P., Rapoport, J., Hoffman, G.E., et al. (2024). Activity-Dependent Transcriptional Program in NGN2+ Neurons Enriched for Genetic Risk for Brain-Related Disorders. Biol. Psychiatry 95, 187–198. 10.1016/j.biopsych.2023.07.003.

12. 12. van Praag, H., Kempermann, G., and Gage, F.H. (2000). Neural consequences of enviromental enrichment. Nat. Rev. Neurosci. 1, 191–198. 10.1038/35044558.

13. Hüttenrauch, M., Salinas, G., and Wirths, O. (2016). Effects of Long-Term Environmental Enrichment on Anxiety, Memory, Hippocampal Plasticity and Overall Brain Gene Expression in C57BL6 Mice. Front. Mol. Neurosci. *9*, 62. 10.3389/fnmol.2016.00062.

14. 14. Dijkhuizen, S., Van Ginneken, L.M.C., IJpelaar, A.H.C., Koekkoek, S.K.E., De Zeeuw, C.I., and Boele, H.J. (2024). Impact of enriched environment on motor performance and learning in mice. Sci. Rep. 14, 5962. 10.1038/s41598-024-56568-3.

15. Escorihuela, R.M., Fernandezteruel, A., Tobena, A., Vivas, N.M., Marmol, F., Badia, A., and Dierssen, M. (1995). Early Environmental Stimulation Produces Long-Lasting Changes on β-Adrenoceptor Transduction System. Neurobiol. Learn. Mem. 64, 49–57. 10.1006/nlme.1995.1043.

16. Leon, M., and Woo, C. (2018). Environmental Enrichment and Successful Aging. Front. Behav. Neurosci. 12. 10.3389/fnbeh.2018.00155.

17. Ball, N.J., Mercado, E., and Orduña, I. (2019). Enriched Environments as a Potential Treatment for Developmental Disorders: A Critical Assessment. Front. Psychol. 10. 10.3389/fpsyg.2019.00466.

18. Nithianantharajah, J., and Hannan, A.J. (2006). Enriched environments, experience-dependent plasticity and disorders of the nervous system. Nat. Rev. Neurosci. 7, 697–709. 10.1038/nrn1970.

19. Krüttner, S., Falasconi, A., Valbuena, S., Galimberti, I., Bouwmeester, T., Arber, S., and Caroni, P. (2022). Absence of familiarity triggers hallmarks of autism in mouse model through aberrant tail-of-striatum and prelimbic cortex signaling. Neuron 110, 1468–1482.e5. 10.1016/j.neuron.2022.02.001.

20. Espeso-Gil, S., Holik, A.Z., Bonnin, S., Jhanwar, S., Chandrasekaran, S., Pique-Regi, R., Albaigès-Ràfols, J., Maher, M., Permanyer, J., Irimia, M., et al. (2021). Environmental Enrichment Induces Epigenomic and Genome Organization Changes Relevant for Cognition. Front. Mol. Neurosci. 14, 664912. 10.3389/fnmol.2021.664912.

21. Rampon, C., Jiang, C.H., Dong, H., Tang, Y.-P., Lockhart, D.J., Schultz, P.G., Tsien, J.Z., and Hu, Y. (2000). Effects of environmental enrichment on gene expression in the brain. Proc. Natl. Acad. Sci. 97, 12880–12884. 10.1073/pnas.97.23.12880.

22. Zhang, T.-Y., Keown, C.L., Wen, X., Li, J., Vousden, D.A., Anacker, C., Bhattacharyya, U., Ryan, R., Diorio, J., O’Toole, N., et al. (2018). Environmental enrichment increases transcriptional and epigenetic differentiation between mouse dorsal and ventral dentate gyrus. Nat. Commun. 9, 298. 10.1038/s41467-017-02748-x.

23. Ali, A.E.A., Wilson, Y.M., and Murphy, M. (2009). A single exposure to an enriched environment stimulates the activation of discrete neuronal populations in the brain of the fos- tau-lacZ mouse. Neurobiol. Learn. Mem. 92, 381–390. 10.1016/j.nlm.2009.05.004.

24. Jaeger, B.N., Linker, S.B., Parylak, S.L., Barron, J.J., Gallina, I.S., Saavedra, C.D., Fitzpatrick, C., Lim, C.K., Schafer, S.T., Lacar, B., et al. (2018). A novel environment-evoked transcriptional signature predicts reactivity in single dentate granule neurons. Nat. Commun. 9, 3084. 10.1038/s41467-018-05418-8.

25. Pollina, E.A., Gilliam, D.T., Landau, A.T., Lin, C., Pajarillo, N., Davis, C.P., Harmin, D.A., Yap, E.-L., Vogel, I.R., Griffith, E.C., et al. (2023). A NPAS4–NuA4 complex couples synaptic activity to DNA repair. Nature 614, 732–741. 10.1038/s41586-023-05711-7.

26. Cembrowski, M.S., Wang, L., Sugino, K., Shields, B.C., and Spruston, N. (2016). Hipposeq: a comprehensive RNA-seq database of gene expression in hippocampal principal neurons. eLife 5, e14997. 10.7554/eLife.14997.

27. 27. Yao, Z., van Velthoven, C.T.J., Kunst, M., Zhang, M., McMillen, D., Lee, C., Jung, W., Goldy, J., Abdelhak, A., Aitken, M., et al. (2023). A high-resolution transcriptomic and spatial atlas of cell types in the whole mouse brain. Nature 624, 317–332. 10.1038/s41586-023-06812-z.

28. 28. Bonnefont, J., Tiberi, L., van den Ameele, J., Potier, D., Gaber, Z.B., Lin, X., Bilheu, A., Herpoel, A., Velez Bravo, F.D., Guillemot, F., et al. (2019). Cortical Neurogenesis Requires Bcl6-Mediated Transcriptional Repression of Multiple Self-Renewal-Promoting Extrinsic Pathways. Neuron 103, 1096–1108.e4. 10.1016/j.neuron.2019.06.027.

29. Vierbuchen, T., Ling, E., Cowley, C.J., Couch, C.H., Wang, X., Harmin, D.A., Roberts, C.W.M., and Greenberg, M.E. (2017). AP-1 Transcription Factors and the BAF Complex Mediate Signal-Dependent Enhancer Selection. Mol. Cell 68, 1067–1082.e12. 10.1016/j.molcel.2017.11.026.

30. Stroud, H., Yang, M.G., Tsitohay, Y.N., Davis, C.P., Sherman, M.A., Hrvatin, S., Ling, E., and Greenberg, M.E. (2020). An Activity-Mediated Transition in Transcription in Early Postnatal Neurons. Neuron 107, 874–890.e8. 10.1016/j.neuron.2020.06.008.

31. 31. Galvan, L., Francelle, L., Gaillard, M.-C., de Longprez, L., Carrillo-de Sauvage, M.-A., Liot, G., Cambon, K., Stimmer, L., Luccantoni, S., Flament, J., et al. (2018). The striatal kinase DCLK3 produces neuroprotection against mutant huntingtin. Brain 141, 1434–1454. 10.1093/brain/awy057.

32. Wallin, R., Cain, D., Hutson, S.M., Sane, D.C., and Loeser, R. (2000). Modulation of the binding of matrix Gla protein (MGP) to bone morphogenetic protein-2 (BMP-2). Thromb. Haemost. 84, 1039–1044.

33. Song, W., Cho, Y., Watt, D., and Cavalli, V. (2015). Tubulin-tyrosine Ligase (TTL)-mediated Increase in Tyrosinated α-Tubulin in Injured Axons Is Required for Retrograde Injury Signaling and Axon Regeneration*. J. Biol. Chem. 290, 14765–14775. 10.1074/jbc.M114.622753.

34. CellTalkDB::About http://tcm.zju.edu.cn/celltalkdb/about.php.

35. Yan, L., Lee, S., Lazzaro, D.R., Aranda, J., Grant, M.B., and Chaqour, B. (2015). Single and Compound Knock-outs of MicroRNA (miRNA)-155 and Its Angiogenic Gene Target *CCN1* in Mice Alter Vascular and Neovascular Growth in the Retina via Resident Microglia*. J. Biol. Chem. 290, 23264–23281. 10.1074/jbc.M115.646950.

36. Microglia in neurodegenerative diseases: mechanism and potential therapeutic targets | Signal Transduction and Targeted Therapy https://www-nature-com.ezp-prod1.hul.harvard.edu/articles/s41392-023-01588-0.

37. Wang, C.S., Kavalali, E.T., and Monteggia, L.M. (2022). BDNF signaling in context: From synaptic regulation to psychiatric disorders. Cell 185, 62–76. 10.1016/j.cell.2021.12.003.

38. Gelfo, F., Mandolesi, L., Serra, L., Sorrentino, G., and Caltagirone, C. (2018). The Neuroprotective Effects of Experience on Cognitive Functions: Evidence from Animal Studies on the Neurobiological Bases of Brain Reserve. Neuroscience 370, 218–235. 10.1016/j.neuroscience.2017.07.065.

39. Wang, Y., Kerrisk Campbell, M., Tom, I., Foreman, O., Hanson, J.E., and Sheng, M. (2020). PCDH7 interacts with GluN1 and regulates dendritic spine morphology and synaptic function. Sci. Rep. 10, 10951. 10.1038/s41598-020-67831-8.

40. Astrotactin 2 (ASTN2) regulates emotional and cognitive functions by affecting neuronal morphogenesis and monoaminergic systems - Ito - 2023 - Journal of Neurochemistry - Wiley Online Library https://onlinelibrary-wiley-com.ezp-prod1.hul.harvard.edu/doi/10.1111/jnc.15790.

41. Hanzel, M., Fernando, K., Maloney, S.E., Gong, S., Mätlik, K., Zhao, J., Pasolli, H.A., Heissel, S., Dougherty, J.D., Hull, C., et al. (2024). Mice lacking Astn2 have ASD-like behaviors and altered cerebellar circuit properties. Preprint at bioRxiv, 10.1101/2024.02.18.580354.

42. Zamora, N.N., Cheli, V.T., González, D.A.S., Wan, R., and Paez, P.M. (2020). Deletion of Voltage-Gated Calcium Channels in Astrocytes during Demyelination Reduces Brain Inflammation and Promotes Myelin Regeneration in Mice. J. Neurosci. 40, 3332–3347. 10.1523/JNEUROSCI.1644-19.2020.

43. Bjursell, M., Ahnmark, A., Bohlooly-Y, M., William-Olsson, L., Rhedin, M., Peng, X.-R., Ploj, K., Gerdin, A.-K., Arnerup, G., Elmgren, A., et al. (2007). Opposing Effects of Adiponectin Receptors 1 and 2 on Energy Metabolism. Diabetes 56, 583–593. 10.2337/db06-1432.

44. Zhang, D., Wang, X., Wang, B., Garza, J.C., Fang, X., Wang, J., Scherer, P.E., Brenner, R., Zhang, W., and Lu, X.-Y. (2017). Adiponectin regulates contextual fear extinction and intrinsic excitability of dentate gyrus granule neurons through AdipoR2 receptors. Mol. Psychiatry 22, 1044–1055. 10.1038/mp.2016.58.

45. Tullai, J.W., Schaffer, M.E., Mullenbrock, S., Sholder, G., Kasif, S., and Cooper, G.M. (2007). Immediate-Early and Delayed Primary Response Genes Are Distinct in Function and Genomic Architecture*. J. Biol. Chem. 282, 23981–23995. 10.1074/jbc.M702044200.

46. Comparative chromatin accessibility upon BDNF stimulation delineates neuronal regulatory elements | Molecular Systems Biology https://www-embopress-org.ezp-prod1.hul.harvard.edu/doi/full/10.15252/msb.202110473.

47. Ichikawa-Tomikawa, N., Ogawa, J., Douet, V., Xu, Z., Kamikubo, Y., Sakurai, T., Kohsaka, S., Chiba, H., Hattori, N., Yamada, Y., et al. (2012). Laminin α1 is essential for mouse cerebellar development. Matrix Biol. 31, 17–28. 10.1016/j.matbio.2011.09.002.

48. Schep, A.N., Wu, B., Buenrostro, J.D., and Greenleaf, W.J. (2017). chromVAR: inferring transcription-factor-associated accessibility from single-cell epigenomic data. Nat. Methods 14, 975–978. 10.1038/nmeth.4401.

49. Zeisel, A., Muñoz-Manchado, A.B., Codeluppi, S., Lönnerberg, P., La Manno, G., Juréus, A., Marques, S., Munguba, H., He, L., Betsholtz, C., et al. (2015). Cell types in the mouse cortex and hippocampus revealed by single-cell RNA-seq. Science 347, 1138–1142. 10.1126/science.aaa1934.

50. Hrvatin, S., Tzeng, C.P., Nagy, M.A., Stroud, H., Koutsioumpa, C., Wilcox, O.F., Assad, E.G., Green, J., Harvey, C.D., Griffith, E.C., et al. (2019). A scalable platform for the development of cell-type-specific viral drivers. eLife 8, e48089. 10.7554/eLife.48089.

51. Vormstein-Schneider, D., Lin, J.D., Pelkey, K.A., Chittajallu, R., Guo, B., Arias-Garcia, M.A., Allaway, K., Sakopoulos, S., Schneider, G., Stevenson, O., et al. (2020). Viral manipulation of functionally distinct interneurons in mice, non-human primates and humans. Nat. Neurosci. 23, 1629–1636. 10.1038/s41593-020-0692-9.

52. Furlanis, E., Dai, M., Garcia, B.L., Vergara, J., Pereira, A., Pelkey, K., Tran, T., Gorissen, B.L., Vlachos, A., Hairston, A., et al. (2024). An enhancer-AAV toolbox to target and manipulate distinct interneuron subtypes. Preprint at bioRxiv, 10.1101/2024.07.17.603924.

53. Yee, C., Xiao, Y., Chen, H., Reddy, A.R., Xu, B., Medwig-Kinney, T.N., Zhang, W., Boyle, A.P., Herbst, W.A., Xiang, Y.K., et al. (2024). An activity-regulated transcriptional program directly drives synaptogenesis. Nat. Neurosci. 27, 1695–1707. 10.1038/s41593-024-01728-x.

54. Kurmangaliyev, Y.Z., Yoo, J., Valdes-Aleman, J., Sanfilippo, P., and Zipursky, S.L. (2020). Transcriptional Programs of Circuit Assembly in the *Drosophila* Visual System. Neuron 108, 1045–1057.e6. 10.1016/j.neuron.2020.10.006.

55. Dong, M., Thennavan, A., Urrutia, E., Li, Y., Perou, C.M., Zou, F., and Jiang, Y. (2020). SCDC: bulk gene expression deconvolution by multiple single-cell RNA sequencing references. Brief. Bioinform. 22, 416–427. 10.1093/bib/bbz166.

56. Kim, D., Paggi, J.M., Park, C., Bennett, C., and Salzberg, S.L. (2019). Graph-based genome alignment and genotyping with HISAT2 and HISAT-genotype. Nat. Biotechnol. 37, 907–915. 10.1038/s41587-019-0201-4.

57. Dobin, A., Davis, C.A., Schlesinger, F., Drenkow, J., Zaleski, C., Jha, S., Batut, P., Chaisson, M., and Gingeras, T.R. (2013). STAR: ultrafast universal RNA-seq aligner. Bioinformatics 29, 15–21. 10.1093/bioinformatics/bts635.

58. featureCounts: an efficient general purpose program for assigning sequence reads to genomic features | Bioinformatics | Oxford Academic https://academic-oup-com.ezp-prod1.hul.harvard.edu/bioinformatics/article/30/7/923/232889.

59. Love, M.I., Huber, W., and Anders, S. (2014). Moderated estimation of fold change and dispersion for RNA-seq data with DESeq2. Genome Biol. 15, 550. 10.1186/s13059-014-0550-8.

60. Zhu, A., Ibrahim, J.G., and Love, M.I. (2019). Heavy-tailed prior distributions for sequence count data: removing the noise and preserving large differences. Bioinformatics 35, 2084– 2092. 10.1093/bioinformatics/bty895.

61. Gu, Z. (2022). Complex heatmap visualization. iMeta 1, e43. 10.1002/imt2.43.

62. 62. Koopmans, F., van Nierop, P., Andres-Alonso, M., Byrnes, A., Cijsouw, T., Coba, M.P., Cornelisse, L.N., Farrell, R.J., Goldschmidt, H.L., Howrigan, D.P., et al. (2019). SynGO: An Evidence-Based, Expert-Curated Knowledge Base for the Synapse. Neuron 103, 217–234.e4. 10.1016/j.neuron.2019.05.002.

63. Zemke, N.R., Armand, E.J., Wang, W., Lee, S., Zhou, J., Li, Y.E., Liu, H., Tian, W., Nery, J.R., Castanon, R.G., et al. (2023). Conserved and divergent gene regulatory programs of the mammalian neocortex. Nature 624, 390–402. 10.1038/s41586-023-06819-6.

64. Wolf, F.A., Angerer, P., and Theis, F.J. (2018). SCANPY: large-scale single-cell gene expression data analysis. Genome Biol. 19, 15. 10.1186/s13059-017-1382-0.

65. Current best practices in single-cell RNA-seq analysis: a tutorial | Molecular Systems Biology https://www-embopress-org.ezp-prod1.hul.harvard.edu/doi/full/10.15252/msb.20188746.

66. Best practices for single-cell analysis across modalities | Nature Reviews Genetics https://www-nature-com.ezp-prod1.hul.harvard.edu/articles/s41576-023-00586-w.

67. Wolock, S.L., Lopez, R., and Klein, A.M. (2019). Scrublet: Computational Identification of Cell Doublets in Single-Cell Transcriptomic Data. Cell Syst. 8, 281–291.e9. 10.1016/j.cels.2018.11.005.

68. Liu, H., Zeng, Q., Zhou, J., Bartlett, A., Wang, B.-A., Berube, P., Tian, W., Kenworthy, M., Altshul, J., Nery, J.R., et al. (2023). Single-cell DNA methylome and 3D multi-omic atlas of the adult mouse brain. Nature 624, 366–377. 10.1038/s41586-023-06805-y.

69. Gayoso, A., Lopez, R., Xing, G., Boyeau, P., Valiollah Pour Amiri, V., Hong, J., Wu, K., Jayasuriya, M., Mehlman, E., Langevin, M., et al. (2022). A Python library for probabilistic analysis of single-cell omics data. Nat. Biotechnol. 40, 163–166. 10.1038/s41587-021-01206-w.

70. McInnes, L., Healy, J., and Melville, J. (2020). UMAP: Uniform Manifold Approximation and Projection for Dimension Reduction. Preprint at arXiv, 10.48550/arXiv.1802.03426.

71. Zhang, K., Zemke, N.R., Armand, E.J., and Ren, B. (2024). A fast, scalable and versatile tool for analysis of single-cell omics data. Nat. Methods 21, 217–227. 10.1038/s41592-023-02139-9.

72. Martens, L.D., Fischer, D.S., Yépez, V.A., Theis, F.J., and Gagneur, J. (2024). Modeling fragment counts improves single-cell ATAC-seq analysis. Nat. Methods 21, 28–31. 10.1038/s41592-023-02112-6.

73. Rauluseviciute, I., Riudavets-Puig, R., Blanc-Mathieu, R., Castro-Mondragon, J.A., Ferenc, K., Kumar, V., Lemma, R.B., Lucas, J., Chèneby, J., Baranasic, D., et al. (2024). JASPAR 2024: 20th anniversary of the open-access database of transcription factor binding profiles. Nucleic Acids Res. 52, D174–D182. 10.1093/nar/gkad1059.

74. Vallat, R. (2018). Pingouin: statistics in Python. J. Open Source Softw. 3, 1026. 10.21105/joss.01026.

75. Heinz, S., Benner, C., Spann, N., Bertolino, E., Lin, Y.C., Laslo, P., Cheng, J.X., Murre, C., Singh, H., and Glass, C.K. (2010). Simple combinations of lineage-determining transcription factors prime cis-regulatory elements required for macrophage and B cell identities. Mol. Cell 38, 576–589. 10.1016/j.molcel.2010.05.004.

