## Supplementary_figures for "Novel environment exposure drives temporally defined and region-specific chromatin accessibility and gene expression changes in the hippocampus"

Figure S1

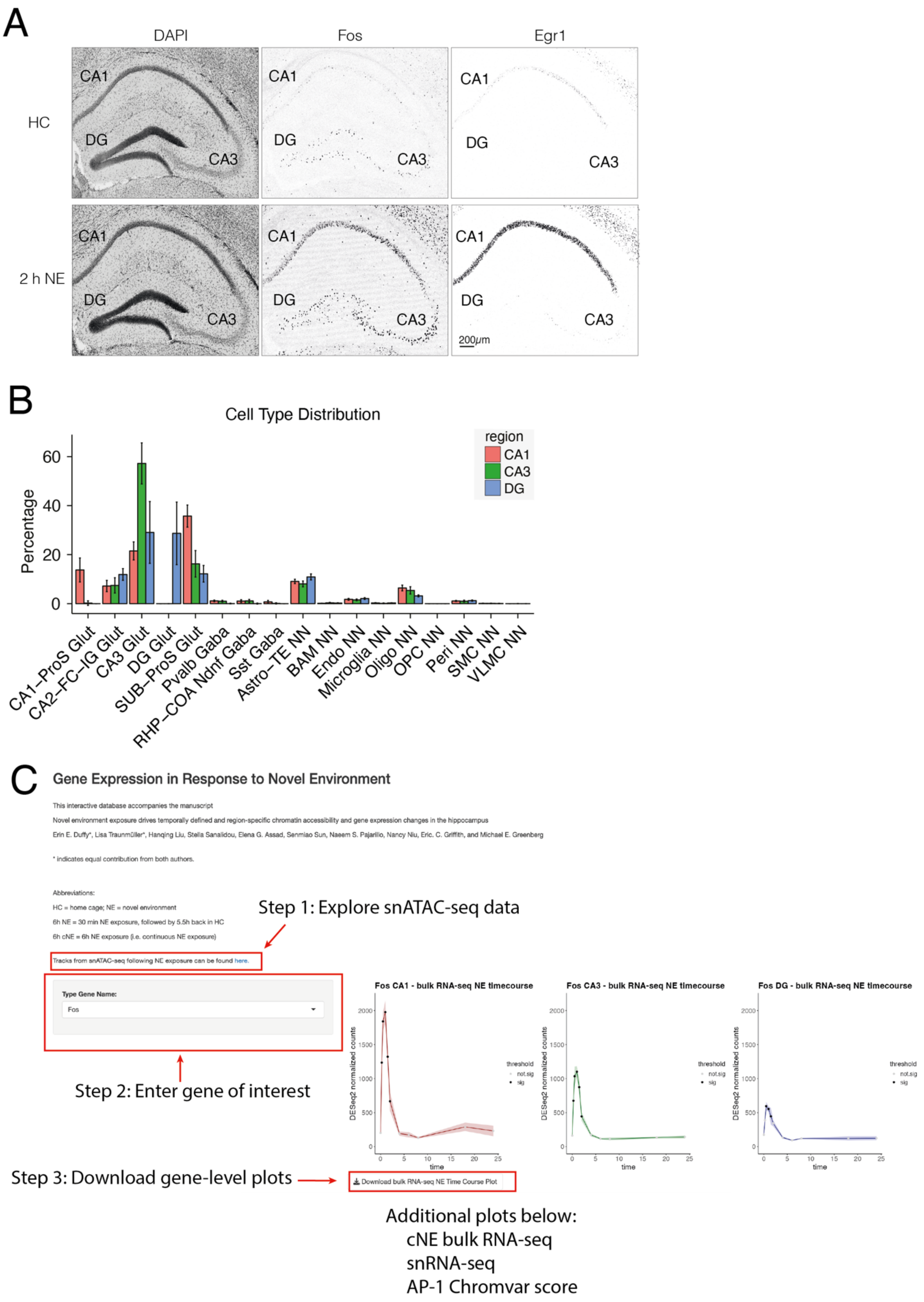

A
DAPI
Fos
Egr1
HC
2 h NE
CA1
DG
CA3
200μm
B
Cell Type Distribution
Percentage
60
40
20
0
region
CA1
CA3
DG
CA1–ProS Glut
CA2–FC–IG Glut
CA3 Glut
DG Glut
SUB–ProS Glut
Pvalb Gaba
RHP–COA Ndnf Gaba
Sst Gaba
Astro–TE NN
BAM NN
Endo NN
Microglia NN
Oligo NN
OPC NN
Peri NN
SMC NN
VLMC NN
C
Gene Expression in Response to Novel Environment
This interactive database accompanies the manuscript
Novel environment exposure drives temporally defined and region-specific chromatin accessibility and gene expression changes in the hippocampus
Erin E. Duffy*, Lisa Traunmüller*, Hanqing Liu, Stella Sanalidou, Elena G. Assad, Senmiao Sun, Naeem S. Pajarillo, Nancy Niu, Eric. C. Griffith, and Michael E. Greenberg
* indicates equal contribution from both authors.
Abbreviations:
HC = home cage; NE = novel environment
6h NE = 30 min NE exposure, followed by 5.5h back in HC
6h cNE = 6h NE exposure (i.e. continuous NE exposure)
Tracks from snATAC-seq following NE exposure can be found here.
Step 1: Explore snATAC-seq data
Type Gene Name:
Fos
Step 2: Enter gene of interest
Step 3: Download gene-level plots
Download bulk RNA-seq NE Time Course Plot
Fos CA1 - bulk RNA-seq NE timecourse
Fos CA3 - bulk RNA-seq NE timecourse
Fos DG - bulk RNA-seq NE timecourse
DESeq2 normalized counts
time
threshold
not.sig
sig
Additional plots below:
cNE bulk RNA-seq
snRNA-seq
AP-1 Chromvar score

**Extended Data Figure 1: NE exposure stimulates hippocampal ERG activity, Related to Figure 1. (A)** Immunofluorescence images of FOS, and EGR1 protein levels in HC and 2 h NE (30 min NE + 1.5 h HC) in CA1, CA3, and DG. Scale bar 200 μm **(B)** As differences in cell type composition can drive differential expression in bulk RNA measurements, we used single-cell deconvolution[55] to approximate the proportion of neuronal and non-neuronal cells in each sample by region, using the Brain Initiative Cell Census Network 2.0[27] as a reference. Bar plot of bulk gene expression deconvolution of RNA-seq from CA1, CA3, and DG across all time points of NE exposure. Differences in non-neuronal cell type composition are not statistically significant by two-way ANOVA (F statistic. = 0.00, P=1.000), strongly suggesting that observed differences between regions are driven primarily by gene expression changes in excitatory cells rather than non-neuronal cells. Data are shown as mean ± SD, n=48-50 biologically independent tissue samples per region. **(C)** Tutorial on using the online database. Users can explore snATAC-seq tracks in the UCSC genome browser, enter a gene of interest, and download high-resolution plots of bulk RNA-seq expression following brief and cNE exposure, as well as cell type-specific gene expression (snRNA-seq) and AP-1 Chromvar scores.

Figure S2

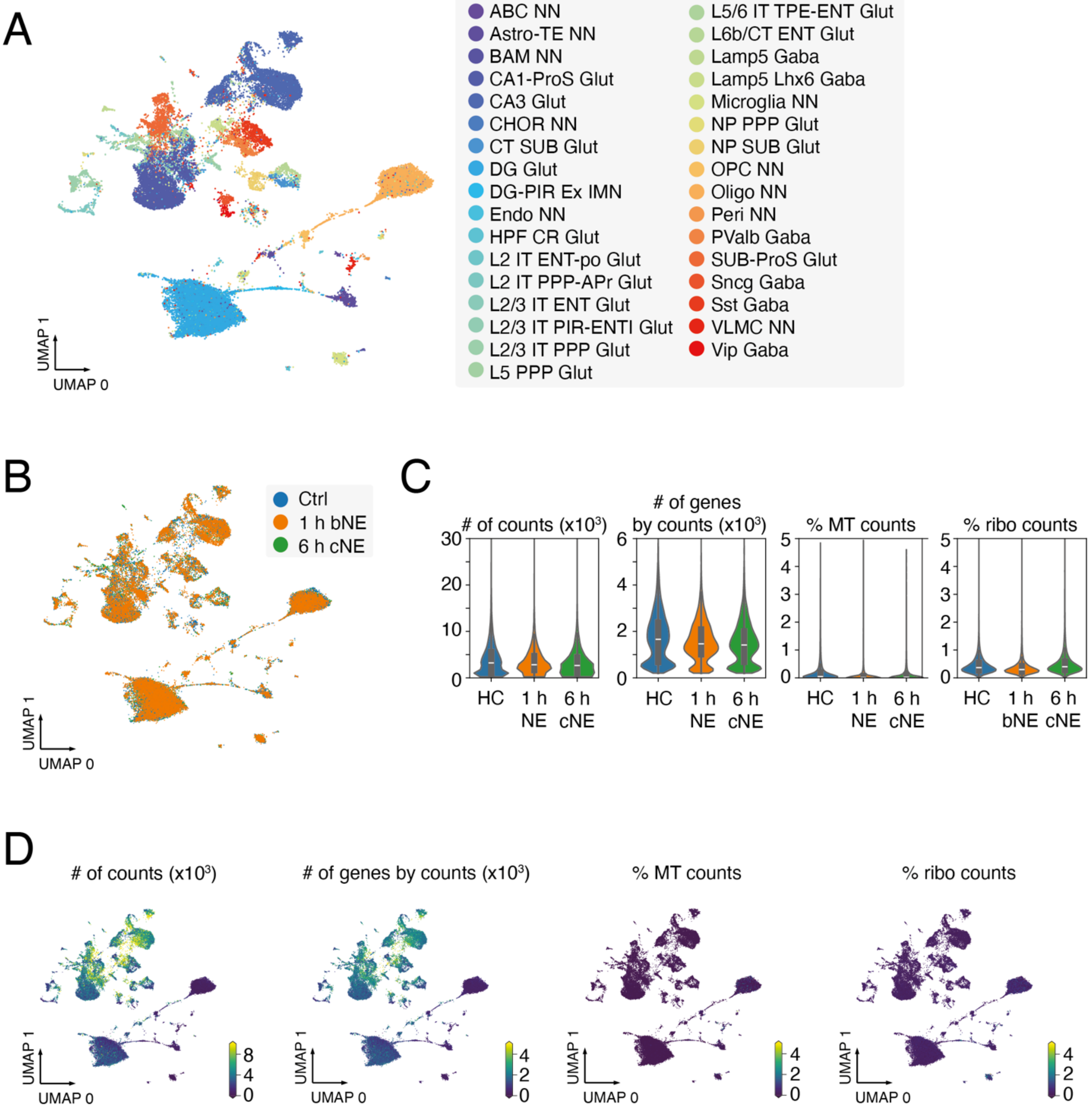


**Extended Data Figure 2: snRNA-seq quality metrics, Related to Figure 1. (A)** UMAP visualization of nuclei from HC, 1 h brief NE (bNE), and 6 h continuous NE (cNE) snRNA-seq with cell sub-type information overlaid. n = 45,844 cells, 3 mice. **(B)** UMAP as in (A), with sample information overlaid. **(C)** Violin plot of number of counts per cell, number of genes by counts, percentage of mitochondrial RNA detected per cell (% MT counts), and percentage of ribosomal RNA detected per cell (%ribo counts), separated by sample (HC, 1 h brief NE, 6 h cNE). **(D)** UMAP as in (A), with information from C overlaid.

### Figure S3

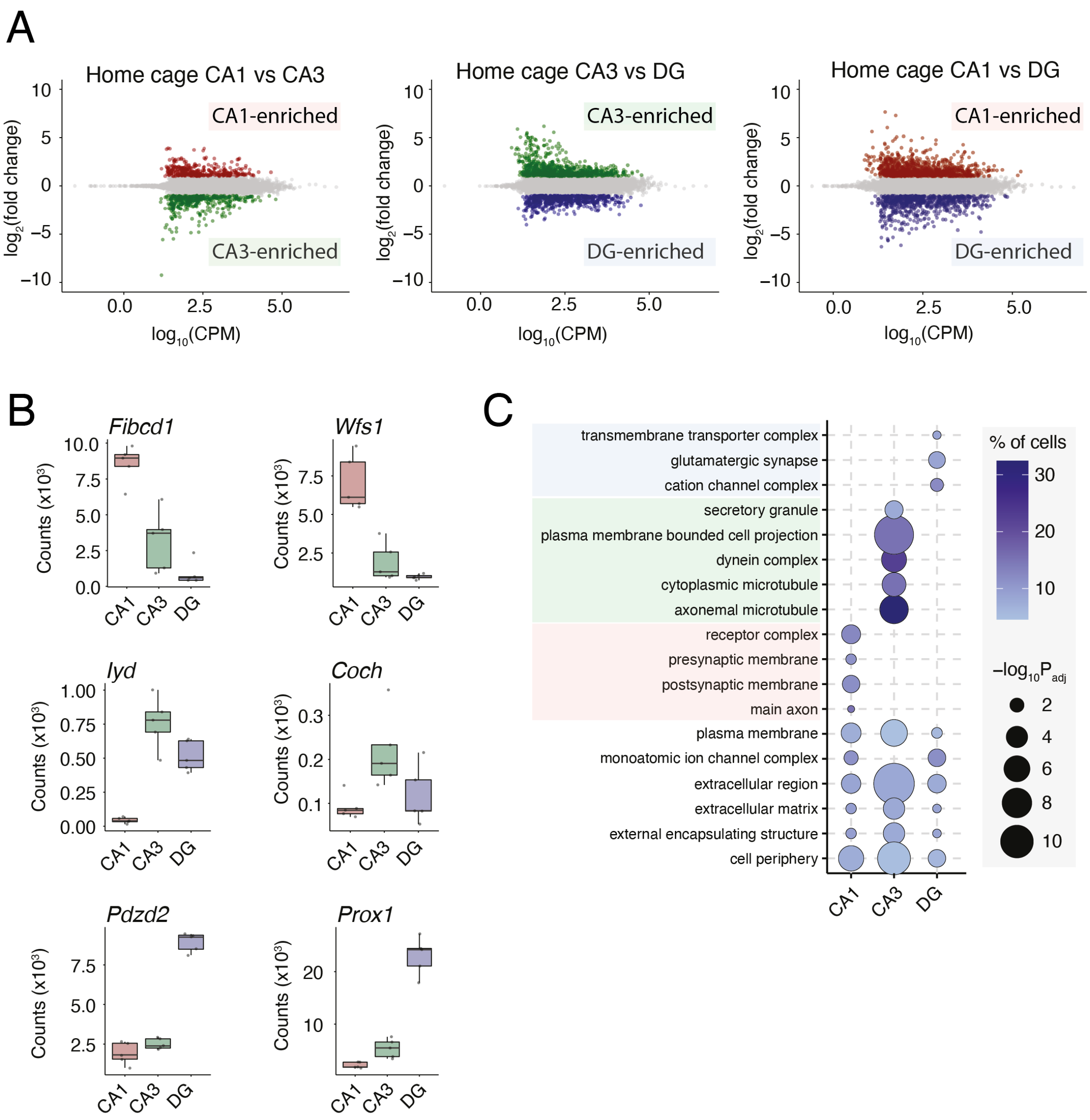


**Extended Data Figure 3: Region-specific differences in HC gene expression, Related to Figure 2. (A)** MA plot of pairwise differentially expressed genes in HC condition between CA1 (red), CA3, (green) and DG (blue). Colored points represent significant (DESeq2 $P_{adj} < 0.05$) enrichment in one region. (**B**) Box and whisker plots of HC bulk RNA-seq depth-normalized counts for genes defined as region-specific marker genes by Cembrowski et al. 2016.[26] **(C)** Dot plot of the top enriched GO terms for CA1- CA3- and DG-enriched genes in Figure 2A, as well as GO terms that were common between all three cell classes.

### Figure S4

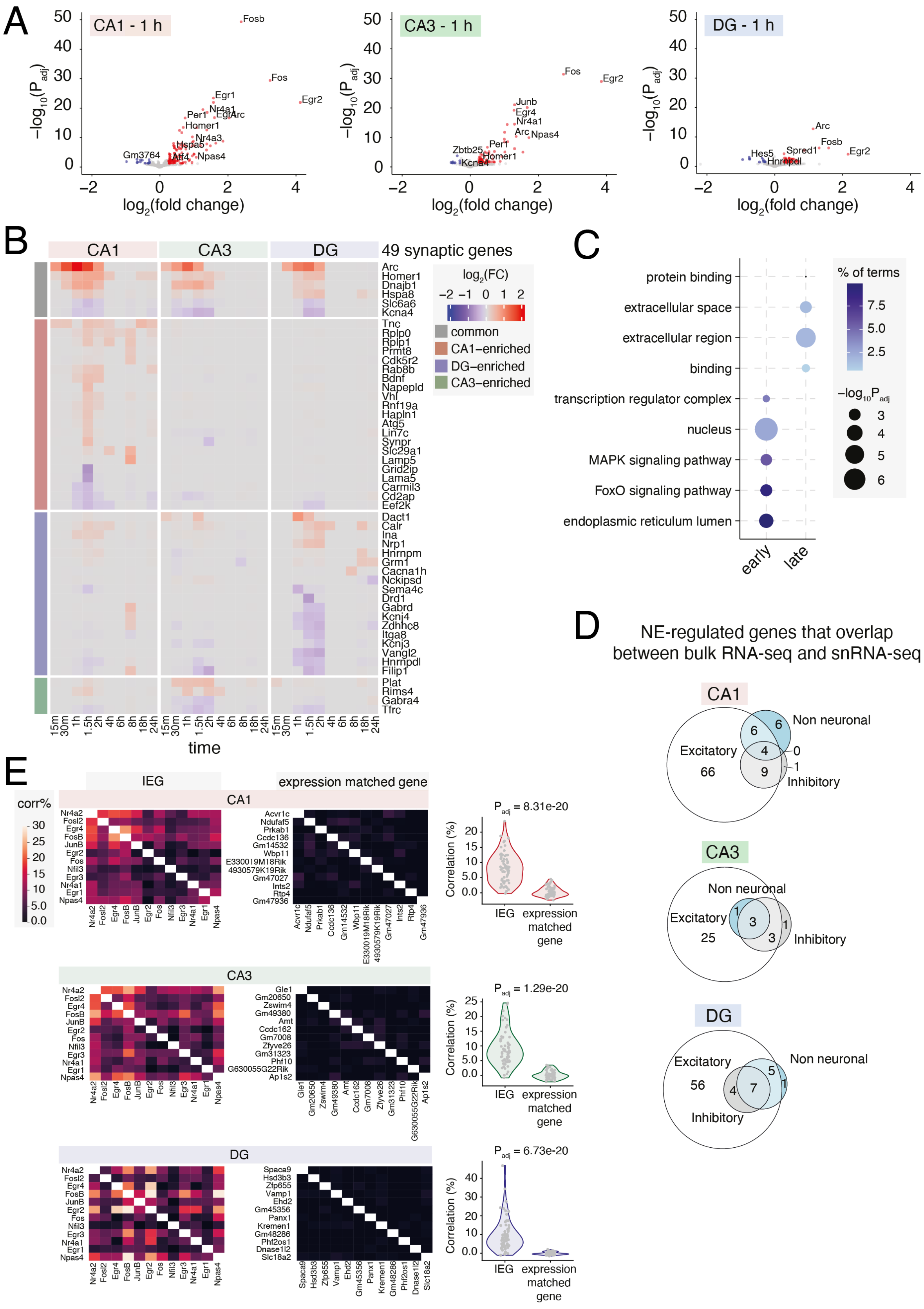

A
CA1 - 1 h
CA3 - 1 h
DG - 1 h
−log10(Padj)
log2(fold change)
Fosb
Fos
Egr2
Egr1
Nr4a1
Per1
Egr4
Arc
Homer1
Nr4a3
Hspa8
Gm3764
Atf4
Npas4
Junb
Zbtb25
Kcna4
Hes5
Spred1
Hnrnpdl
B
CA1
CA3
DG
49 synaptic genes
log2(FC)
common
CA1−enriched
DG−enriched
CA3−enriched
Arc
Homer1
Dnajb1
Hspa8
Slc6a6
Kcna4
Tnc
Rplp0
Rplp1
Prmt8
Cdk5r2
Rab8b
Bdnf
Napepld
Vhl
Rnf19a
Hapln1
Atg5
Lin7c
Synpr
Slc29a1
Lamp5
Grid2ip
Lama5
Carmil3
Cd2ap
Eef2k
Dact1
Calr
Ina
Nrp1
Hnrnpm
Grm1
Cacna1h
Nckipsd
Sema4c
Drd1
Gabrd
Kcnj4
Zdhhc8
Itga8
Kcnj3
Vangl2
Hnrnpdl
Filip1
Plat
Rims4
Gabra4
Tfrc
15m 30m 1h 1.5h 2h 4h 6h 8h 18h 24h
time
C
protein binding
extracellular space
extracellular region
binding
transcription regulator complex
nucleus
MAPK signaling pathway
FoxO signaling pathway
endoplasmic reticulum lumen
early
late
% of terms
7.5
5.0
2.5
−log10Padj
3
4
5
6
D
NE-regulated genes that overlap between bulk RNA-seq and snRNA-seq
CA1
Non neuronal
Excitatory
66
Inhibitory
CA3
Non neuronal
Excitatory
25
Inhibitory
DG
Excitatory
56
Non neuronal
Inhibitory
E
IEG
expression matched gene
corr%
CA1
CA3
DG
Nr4a2
Fosl2
Egr4
FosB
JunB
Egr2
Fos
Nfil3
Egr3
Nr4a1
Egr1
Npas4
Acvr1c
Ndufaf5
Prkab1
Ccdc136
Gm14532
Wbp11
E330019M18Rik
4930579K19Rik
Gm47027
Ints2
Rtp4
Gm47936
Gle1
Gm20650
Zswim4
Gm49380
Amt
Ccdc162
Gm7008
Zfyve26
Gm31323
Phf10
G630055G22Rik
Ap1s2
Spaca9
Hsd3b3
Zfp655
Vamp1
Ehd2
Gm45356
Panx1
Kremen1
Gm48286
Phf2os1
Dnase1l2
Slc18a2
Correlation (%)
Padj = 8.31e-20
Padj = 1.29e-20
Padj = 6.73e-20
IEG
expression matched gene

**Extended Data Figure 4: NE-driven gene expression changes, Related to Figure 3. (A)** Volcano plot of $-\log_{10}(P_{adj})$ versus $\log_2$(fold change) in bulk RNA-seq expression between 1 h brief NE exposure and HC in CA1, CA3, and DG. Red indicates upregulated genes (DEseq2 $P_{adj} < 0.05$, $\text{Log}_2$ fold change $> \log_2(1.2)$); blue indicates downregulated genes (DEseq2 $P_{adj} < 0.05$, $\text{Log}_2$ fold change $< -\log_2(1.2)$); and gray indicates non-significant genes (DEseq2 $P_{adj} > 0.05$). **(B)** Heatmap as in (Fig 3C) for genes that are annotated as synaptic genes. **(C)** Dot plot of the top GO terms in early and late upregulated genes by bulk RNA-seq. ERGs were enriched for GO terms related to transcription factor activity in the nucleus, whereas late response genes were enriched for terms related to extracellular secretion and receptor binding. **(D)** Venn diagrams of genes detected as differentially expressed in both bulk and snRNA-seq. Genes were classified as CA1 CA3 or DG based on differential expression in bulk RNA-seq and further classified as excitatory, inhibitory, or non-neuronal based on the cell type(s) in which the same gene was differentially expressed in snRNA-seq data at 1 h NE. **(E)** Pairwise Pearson correlations across individual excitatory neurons in CA1, CA3, or DG at 1 h NE exposure, calculated on the basis of (left) commonly induced TF expression (**Fig. 3D**), or (right) expression-matched non-induced genes. Correlations between ERG TFs are significantly higher than those between expression-matched noninduced genes (p-values as indicated, Mann-Whitney *U* test, two-sided).

### Figure S5

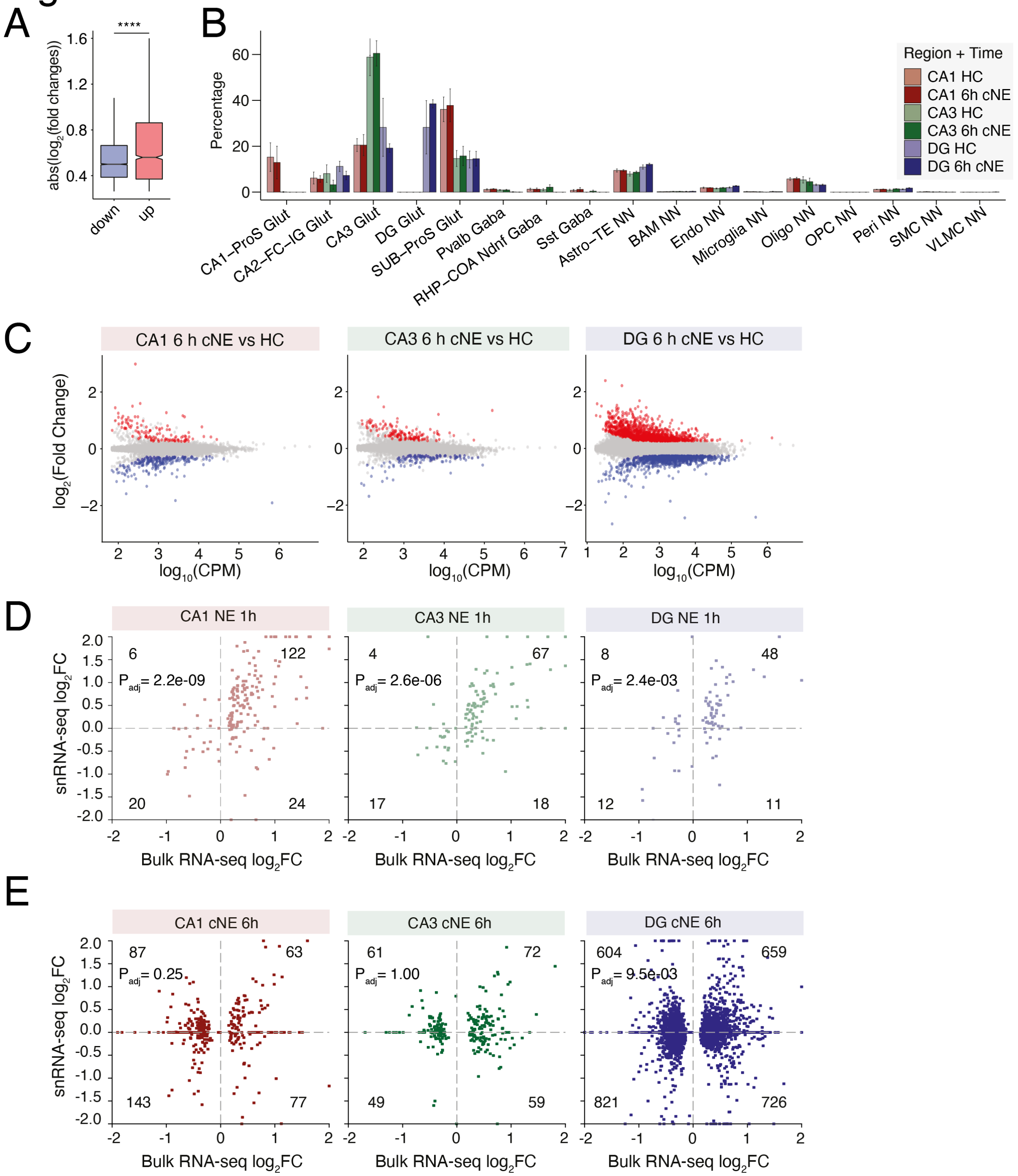


**Extended Data Figure 5: Continuous NE exposure boosts expression of LRGs, Related to Figure 6. (A)** Box plot of absolute value($\log_2$ fold changes) in NE-upregulated (red) and -downregulated (blue) genes by snRNA-seq in Figure 5A. **** = $P < 2.2e\text{-}16$, Welch's Two Sample t-test. **(B)** Bar plot of bulk gene expression deconvolution of RNA-seq from CA1, CA3, and DG, separated by HC and cNE exposure, using the Mouse Brain Atlas as a reference. Differences in cell type composition are not statistically significant between HC and cNE for each region by two-way ANOVA (F statistic = 0.048, P-value = 1.000), strongly suggesting that observed gene expression changes are driven by exposure to cNE and not cell type differences. Data are shown as mean ± SD, n=4-5 biologically independent replicates per region and time. **(C)** MA plot of pairwise differentially expressed genes between HC and 6 h cNE in CA1, CA3, and DG. Red indicates upregulated genes (DEseq2 $P_{adj} < 0.05$, FC > 1.5); blue indicates downregulated genes (DEseq2 $P_{adj} < 0.05$, FC < 0.667); and gray indicates non-significant genes (DEseq2 $P_{adj} > 0.05$). **(D & E)** Scatterplot of $log_2FC$ between bulk RNA-seq (x-axis) and snRNA-seq (y-axis) for genes defined as significant (DESeq2 Padj < 0.05) in bulk RNA-seq within region (CA1, CA3, DG) and time point (D = 1 h brief NE, E = 6 h cNE). $P_{adj}$ = Pearson's R correlation.

### Figure S6

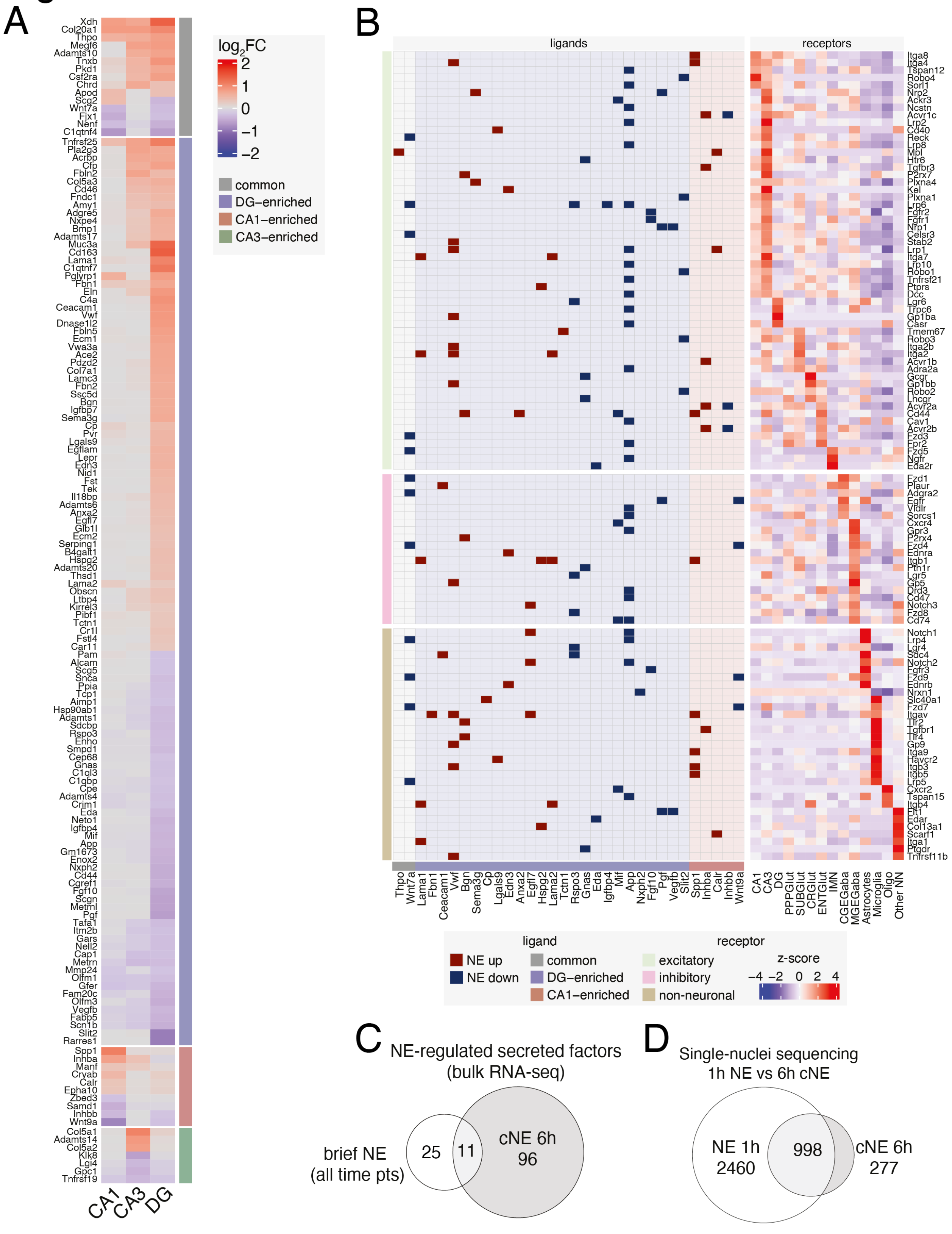

A
log2FC
common
DG–enriched
CA1–enriched
CA3–enriched
CA1
CA3
DG
B
ligands
receptors
ligand
NE up
NE down
common
DG–enriched
CA1–enriched
receptor
excitatory
inhibitory
non-neuronal
z-score
C
NE-regulated secreted factors
(bulk RNA-seq)
brief NE
(all time pts)
25
11
cNE 6h
96
D
Single-nuclei sequencing
1h NE vs 6h cNE
NE 1h
2460
998
cNE 6h
277

**Extended Data Figure 6: Secreted factors induced by 6 h cNE and their receptors, Related to Figure 6.** **(A)** Heatmap of $log_2$(fold change) in gene expression of all significant NE-regulated genes that are annotated secreted factors at 6 h cNE versus HC by bulk RNA-seq. Colors over gene names indicate that the gene has at least one known receptor. **(B)** Interaction grid of NE-regulated secreted factors (ligands) and their receptors. Left grid shows ligand-receptor pair, with dark shading indicating whether ligand expression was upregulated (dark red) or downregulated (dark blue) in response to NE. Background shading indicates the hippocampal region of ligand NE gene expression regulation (common, CA1- CA3- or DG-enriched). Right heatmap shows row-normalized z-scores of cognate receptor expression from HC snRNA-seq across all major cell types, ordered by cell type of maximum receptor expression. (**C**) Venn diagram of secreted factors that are NE-induced following brief (all time points) or 6 h cNE exposure as displayed in Fig4C and S6A, respectively. (**D**) Venn diagram of all differentially regulated genes in snRNA-seq between 1 h brief and 6 h cNE.

### Figure S7

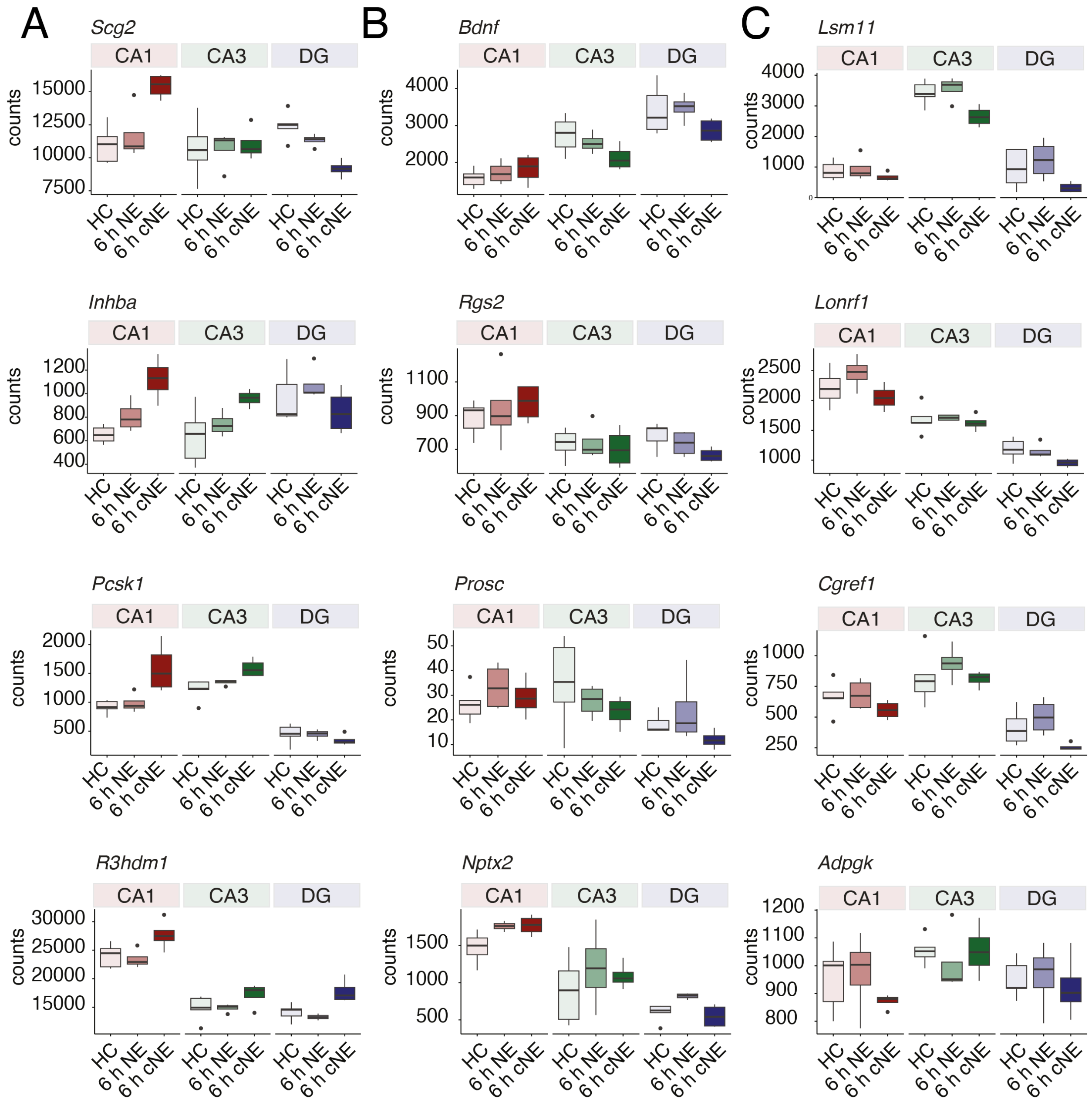


**Extended Data Figure 7: Gene expression changes of AP-1 targets following NE exposure, Related to Figure 6.** Box and whisker plots of bulk RNA-seq depth-normalized counts of gene expression in HC and following brief and cNE exposure for genes previously reported to be activated by Fos/AP-1 in response to kainic acid-induced seizures in CA1.[6] Box plots shown as median ± IQR (whiskers = 1.5*IQR).

### Figure S8

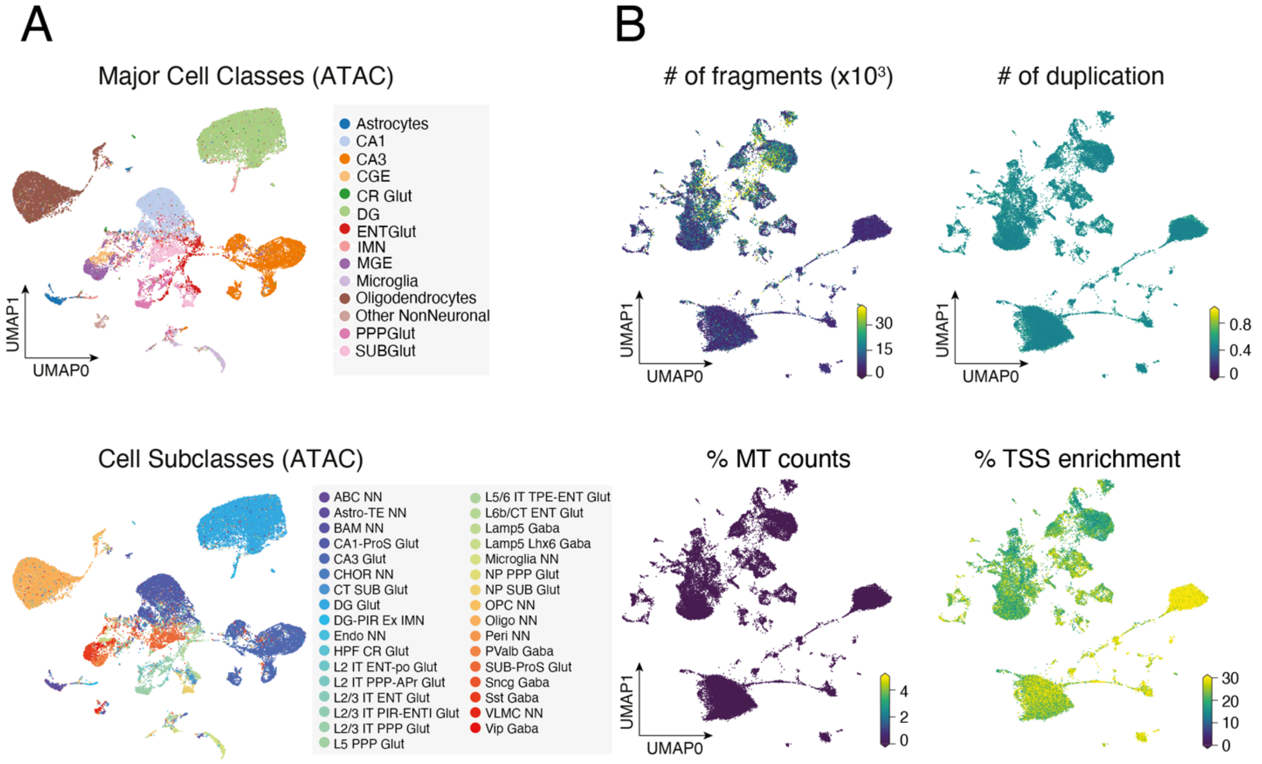


**Extended Data Figure 8: Chromatin accessibility changes in response to NE at single-cell resolution, Related to Figure 7. (A)** UMAP visualization of nuclei from HC, 1 h brief NE, and 6 h cNE snATAC-seq with cell type (top) and cell sub-type (bottom) information overlaid. n = 45,844 cells, 3 mice. **(B)** UMAP as in (Fig 1C), with number of fragments per cell, fraction of duplicated fragments per cell (# of duplication), fraction of mitochondrial fragments per cell (% MT counts), and % transcription start site enrichment (% TSS enrichment) information overlaid.

### Figure S9

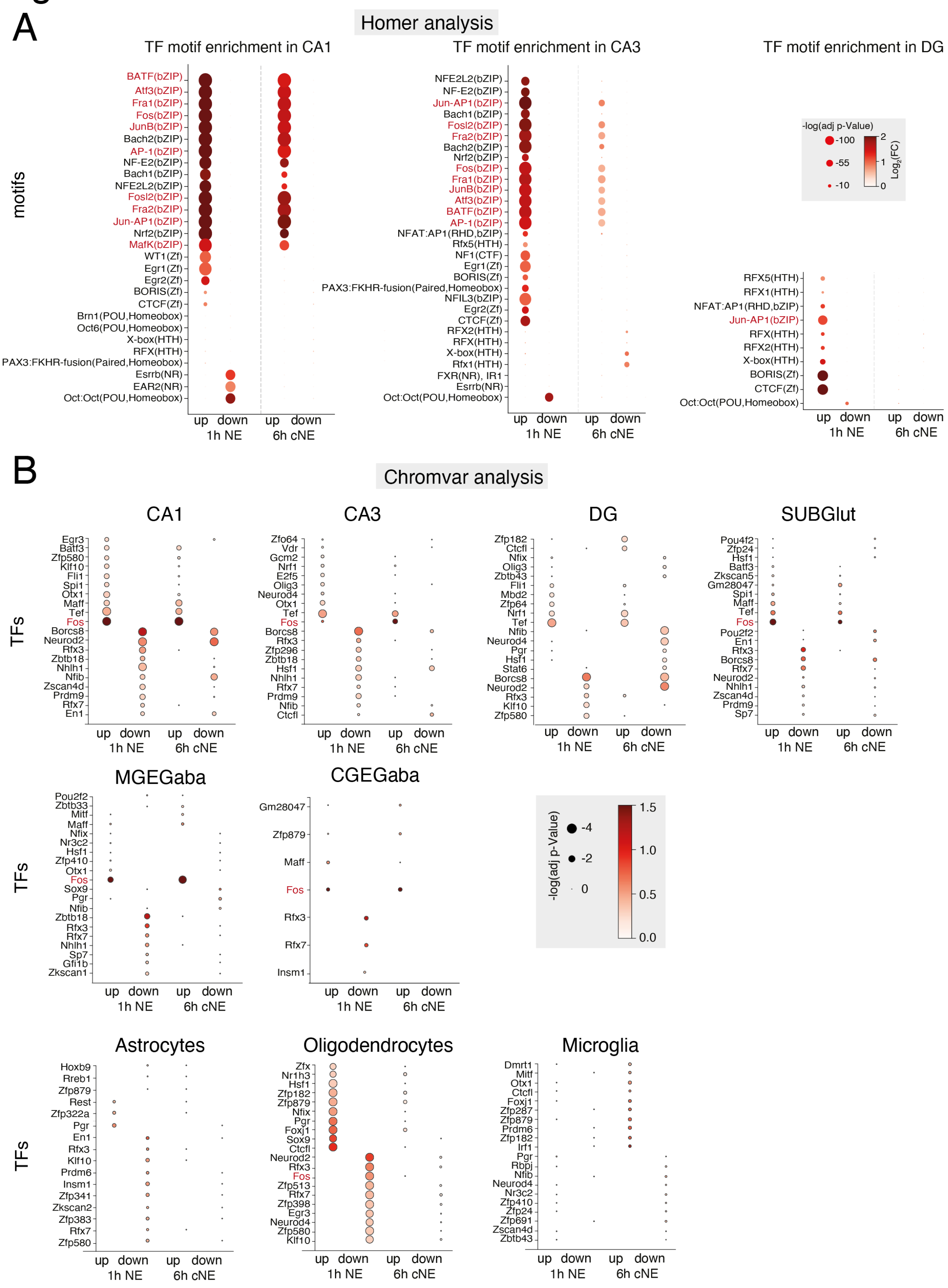


**Extended Data Figure 9: Motif enrichment in activity-regulated ATAC peaks, Related to Figure 7. (A)** Dot plot of TF motifs enriched in CA1, CA3, and DG excitatory neurons within differentially accessible peaks following 1 h brief NE or 6 h cNE compared to HC using Homer.

AP-1-related motifs are indicated in red. **(B)** Dot plot of TF motifs enriched in major excitatory- and inhibitory neurons, and non-neuronal cells within differentially accessible peaks following 1 h brief NE or 6 h cNE compared to HC using Chromvar. Fos is indicated in red.

Figure S10

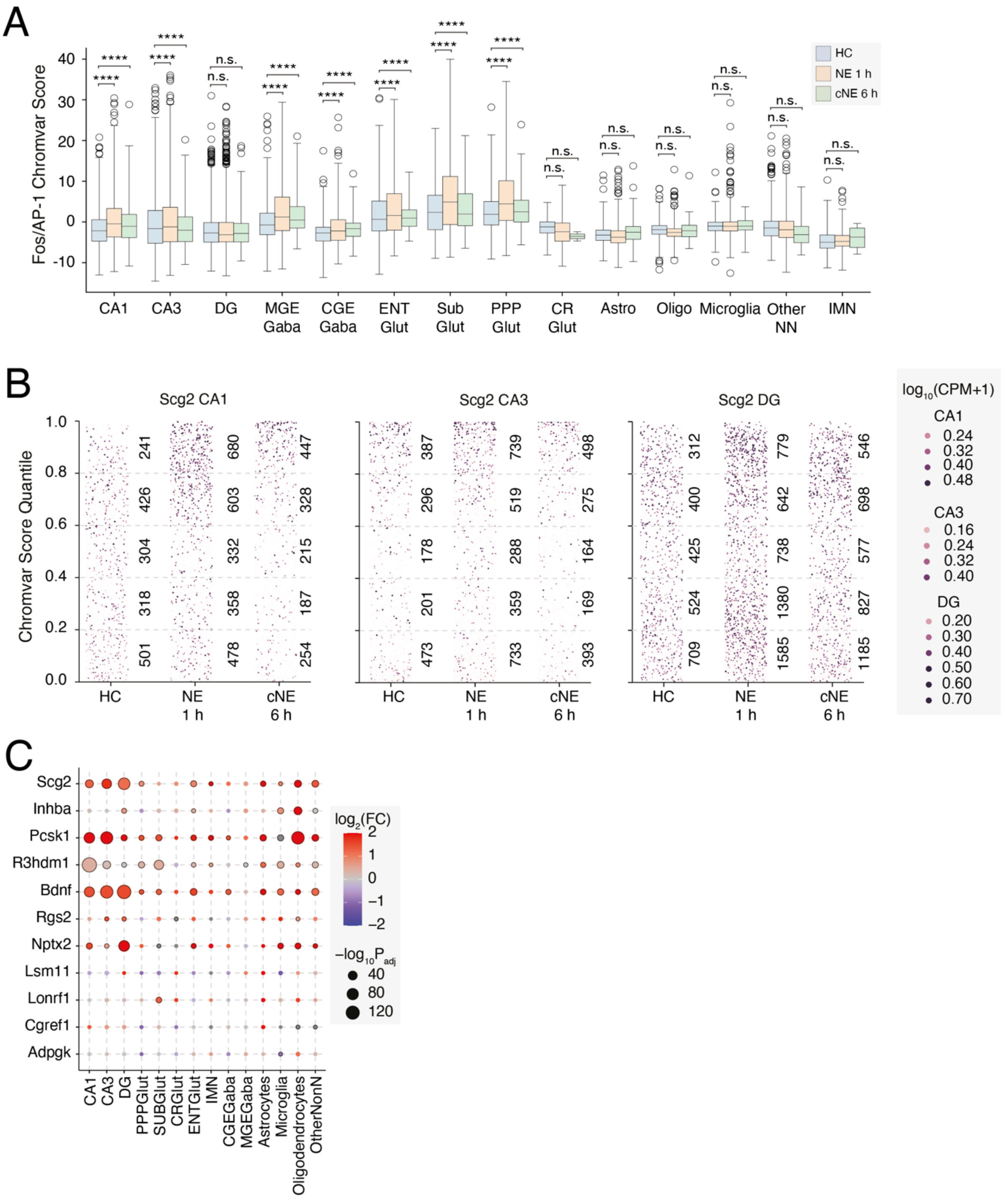


**Extended Data Figure 10: Motif enrichment in activity-regulated ATAC peaks, Related to Figure 7. (A)** Box and whisker plot of Fos/AP-1 Chromvar scores within all major cell types. (**B**) Dot plot of *Scg2* expression in CA1, CA3, and DG excitatory neurons. Cells are plotted based on Fos/AP-1 Chromvar score, where dotted lines indicate Chromvar score quantiles, numbers indicate the number of cells in each quantile, and color indicates *Scg2* expression in $\log_{10}$CPM. **(C)** Dot plot of $\log_2$FC in Fos/AP-1 Chromvar scores of genes displaying up-regulation in bulk RNA-seq after 6 h cNE, no change or down-regulation based on Fig.S7.

**TABLES**

**Table 1:** Fold changes and adjusted p-values for all comparisons in bulk and snRNA-seq
**Table 2:** GO analysis enriched terms
**Table 3:** Chromvar scores for each gene and cell type
